# Enzyme Intermediates Captured “on-the-fly” by Mix-and-Inject Serial Crystallography

**DOI:** 10.1101/202432

**Authors:** Jose Olmos, Suraj Pandey, Jose M. Martin-Garcia, George Calvey, Andrea Katz, Juraj Knoska, Christopher Kupitz, Mark S. Hunter, Mengning Liang, Dominik Oberthuer, Oleksandr Yefanov, Max Wiedorn, Michael Heyman, Mark Holl, Kanupriya Pande, Anton Barty, Mitchell D. Miller, Stephan Stern, Shatabdi Roy-Chowdhury, Jesse Coe, Nirupa Nagaratnam, James Zook, Jacob Verburgt, Tyler Norwood, Ishwor Poudyal, David Xu, Jason Koglin, Matt Seaberg, Yun Zhao, Saša Bajt, Thomas Grant, Valerio Mariani, Garrett Nelson, Ganesh Subramanian, Euiyoung Bae, Raimund Fromme, Russel Fung, Peter Schwander, Matthias Frank, Thomas White, Uwe Weierstall, Nadia Zatsepin, John Spence, Petra Fromme, Henry N. Chapman, Lois Pollack, Lee Tremblay, Abbas Ourmazd, George N. Phillips, Marius Schmidt

## Abstract

Ever since the first atomic structure of an enzyme was solved, the discovery of the mechanism and dynamics of reactions catalyzed by biomolecules has been the key goal for the understanding of the molecular processes that drive life on earth. Despite a large number of successful methods for trapping reaction intermediates, the direct observation of an ongoing reaction has been possible only in rare and exceptional cases. Here, we demonstrate a general method for capturing enzyme catalysis ‘in-action’ by ‘mix-and-inject serial crystallography’. Specifically, we follow the catalytic reaction of the *Mycobacterium tuberculosis* β-lactamase with the 3^rd^ generation antibiotic ceftriaxone by time-resolved serial femtosecond crystallography. The results reveal, in near atomic detail, antibiotic cleavage and inactivation on the millisecond to second time scales including the crossover from transition state kinetics to steady-state kinetics.

**Synopsis:** An enzymatically catalyzed reaction is initiated by diffusion based mixing of substrate and followed at runtime by time-resolved serial crystallography using a free electron laser.

---

Observing the catalytic action of a biomolecule in atomic detail has been the dream of structural biologists since the first structure of an enzyme was solved (*1*, *2*). By exploiting X-ray radiation from powerful synchrotron sources, time-resolved crystallographic methods were developed (*3*) with the goal to achieve a complete description of a reaction in real time (*4*, *5*). However, X-ray damage and the need for large single crystals made time-resolved crystallography very challenging. The advent of X-ray Free Electron Lasers (XFELs) enabled time resolved serial femtosecond (fs) crystallography, where X-ray damage is outrun by ultrashort fs X-ray pulses (*6*, *7*). This approach has made it possible to follow and describe cyclic and non-cyclic reactions that can be triggered by light. Examples include pioneering studies that investigate the photocycle in the photactive yellow protein (*8*, *9*), myoglobin (*10*), and photosystem II (*11*-*14*). However, structural investigations on one-pathway enzymatic reactions present additional difficulties, because diffusion of substrate(s) and products in and out of the crystals limit the accessible reaction times. Standard crystallography can be used to track reaction intermediates of slow reactions by flash freezing but is then unable to reveal enzymatic reactions at room temperature in real time. The problem is to start a reaction in large-sized crystals. Initiation by diffusion is far slower in these crystals than the typical millisecond turnover times of enzymes. It was proposed that one can trigger enzymatic reactions by light by soaking inactive (caged) substrates (*15*) into the crystals, which can be activated for example by a laser pulse. The first proof of concept for TR-Laue crystallography triggered by a caged substrate was achieved in 1990 (*16*). While this method has great potential, its application has so far been limited due to significant experimental challenges. Only a few time-resolved experiments have been reported where highly reactive, caged substrates are readily available (*16*-*18*), or the reactions are slow to allow the use of more conventional methods (*19*, *20*). It is therefore highly desirable to develop new methods that open the field of time-resolved crystallography to the study of biomolecular reactions at room temperature with the native enzyme and its natural substrate(s). Structural studies at XFELs provide a breakthrough. The XFEL intensity is high enough to generate a diffraction pattern from an exposure to a single fs X-ray pulse even from micron and submicron sized crystals. These tiny crystals allow for fast (sub-ms to ms) diffusion times which are not rate limiting for many enzymatic reactions (*21*-*26*). The nanocrystals are mixed “on the fly” and injected into the XFEL beam, a method we call “mix-and-inject serial crystallography” (MISC) (*23*, *25*). In MISC, crystals react with their native substrate(s) at ambient temperature until they are intercepted (probed) the by a single X-ray pulse, which destroys them only after diffraction data has been recorded. The pulses are short enough to essentially outrun radiation damage by means of the “diffraction-before-destruction” principle (*27*-*29*). Optimized injectors have been recently developed (*30*, *31*) for MISC experiments with the potential to provide submillisecond time resolution (*32*). The microcrystals tolerate even larger conformational changes leading to unit cell or even space group changes (*11*, *26*).

Here, we apply MISC to the study of a very important public-health problem: antibiotic resistance of bacteria. Specifically, we have obtained time-resolved crystallographic data on the binding and cleavage of the third-generation antibiotic ceftriaxone (CEF) in microcrystals of the enzyme β-lactamase from *M. tuberculosis* (BlaC). In these experiments carried out at the Linac Coherent Light Source (LCLS) BlaC micro-crystals are mixed with CEF on the fly, and the cleavage and thereby inactivation of the antibiotics by β-lactamase is followed in runtime. BlaC is a broad-spectrum β-lactamase which provides tuberculosis with resistance to all classes of β-lactam antibiotics. BlaC chemistry has rendered the frontline arsenal of antibacterial agents ineffective against this deadly disease, creating a global public health crisis. More generally, our approach is applicable to broad classes of important enzymes with the potential to fundamentally alter our understanding of the molecular basis of biomolecular reactions vital to the design of novel drugs.

Beginning with the famous discovery of penicillin, β-lactam antibiotics were widely used to eliminate deadly infectious diseases (*33*). More compounds with diverse chemical composition were found through the years (*34*), the most prominent of them are most likely the cephalosporins. The chemical structure of CEF is shown in Scheme 1. Unlike the penicillins which feature a 5-membered thiazolidine ring, in the chephalosporins a 6-membered dihydrothiazine ring is fused to the β-lactam ring. However, resistance against these antibiotics was observed shortly after their widespread use, and is now rampant. β-lactamases open the β-lactam ring rendering the antibiotic inactive. BlaC from *M. tuberculosis*, an Ambler class A β-lactamase (*35*), uses a conserved serine to attack the β-lactam ring (scheme 1, blue arrow) thus inactivating the antibiotics. Because of the great medical challenge that BlaC causes for the fight against infectious diseases the process of catalysis has been studied by conventional biochemcial methods in detail leading to the hypothesis of a three step model of the cleavage process: The first step is the formation of the enzyme-substrate complex (species 1), and it has been proposed that the enzyme may use active site interactions to orient the β-lactam carbonyl-carbon near the Ser-70 nucleophile (*36*, *37*). The next step proposed along the reaction coordinate is the nucleophilic attack of Ser-70, which results in the opening of the β-lactam ring and the formation of the covalently bound active site acyl-intermediate (species 3). For cephalosporins there is evidence that during the enzymatic reaction a leaving group (denoted R in Scheme 1) is split off (*38*, *39*). In the third step, the open-ring β-lactam ligand is hydrolyzed and is released by the enzyme (not shown in scheme 1). Various rates have been reported for this step of the catalytic reaction across different classes of β-lactams, followed by product release (*36*). Obtaining time-resolved data on BlaC chemistry holds the potential to directly visualize substrate chemical intermediates and the accompanying active site interactions, for wide-ranging implications for all classes of β-lactams. Ultimately, knowledge of the physical processes by which BlaC is able to bind and catalyze the breakdown of β-lactams, will directly impact rational drug design against deadly human diseases.

**Scheme 1.**
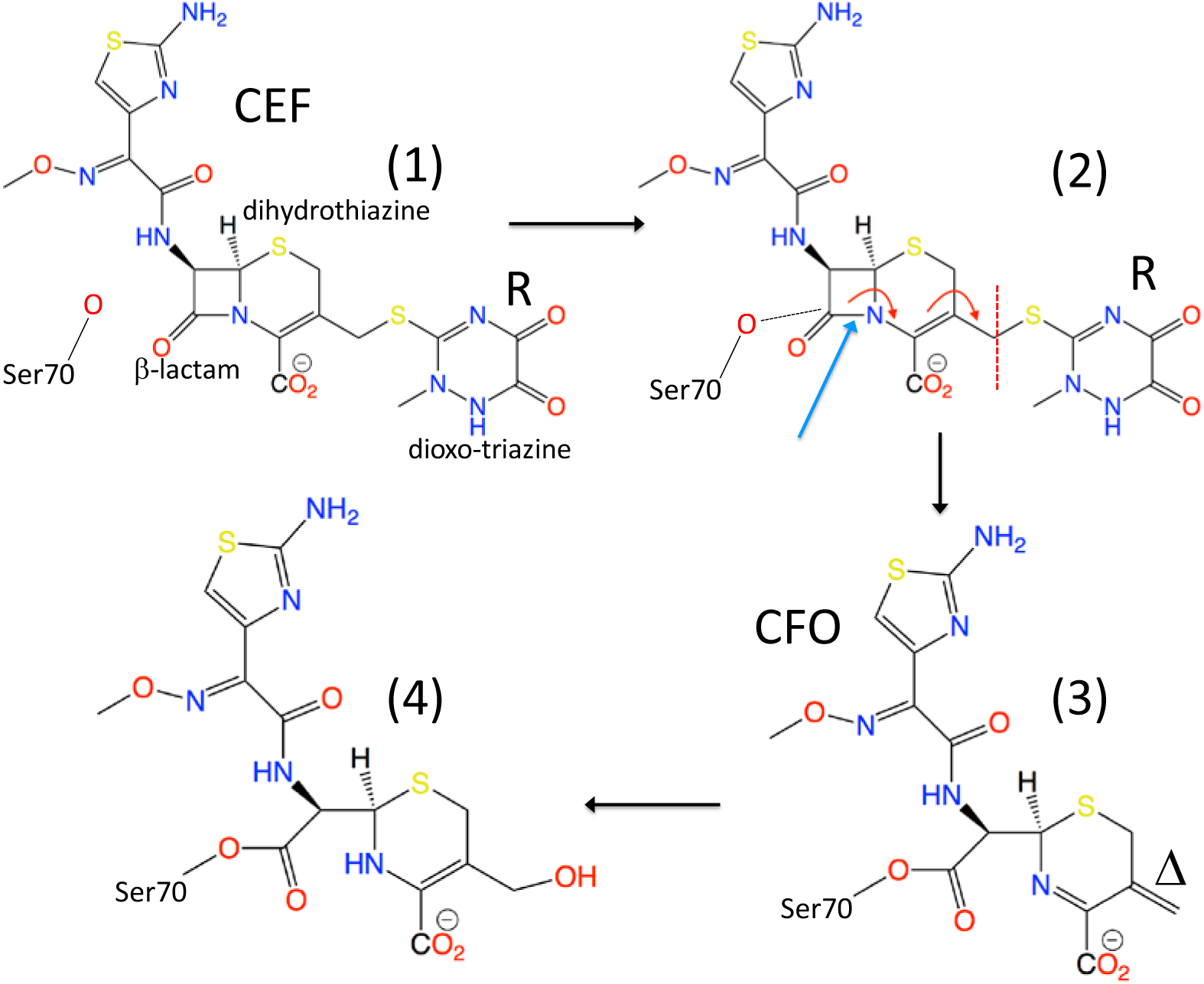
Reaction of β-lactamase with ceftriaxone (CEF). **(1)** Formation of the enzyme substrate complex by non-covalently binding CEF. **(2)** Nucleophilic attack of the active site residue Ser-70 results in rearrangement of double bonds, and ultimately leads to the opening of the β-lactam ring (blue arrow points to the bond to be cleaved), and the detachment of the leaving group (R). **(3)** Covalent bond formation between Ser-70 and a shortened species (E-CFO*). Note the double bond Δ. **(4)** The double bond may react with water to form an alcohol (OH). Evidence for all four intermediate species is found in our experiments. Species (4) is further hydrolyzed from Ser-70 and leaves the enzyme as product.

Our previous results showed that CEF can diffuse into the crystals and binds to the active site of the crystalline tetrameric β-lactamase (*25*). These first studies showed that the catalytic reaction is heterogeneous as the reactivity is specific to individual copies of the four β-lactamase chains in the asymmetric unit of the crystal. Only subunits B and D bind and process CEF, while subunits A and C do not directly contribute to catalysis, as least on the time scale of our experiments (Fig. 1a). This first proof of concept study was limited to a single time point about 2 s after reaction initiation. Multiple time points that cover the reaction are required for any kinetic analysis. Here we present time series from 30 ms to 2 s after mixing with substrate in two crystal forms, shards and needles, which allow us to discover the conformational changes and to characterize the kinetics of this important class of enzymes directly from the X-ray data. We calculate simulated annealing (SA) omit maps at each time point of the time series, and base our interpretation on these bias-free maps. Fig. 2, and supplemental material (SM) Figs. S2 – S6 show details of these maps near the active site. As a complement, more conventional 2mFo-Fc maps are shown in Figs. S7 – S9.

**Figure 1.**
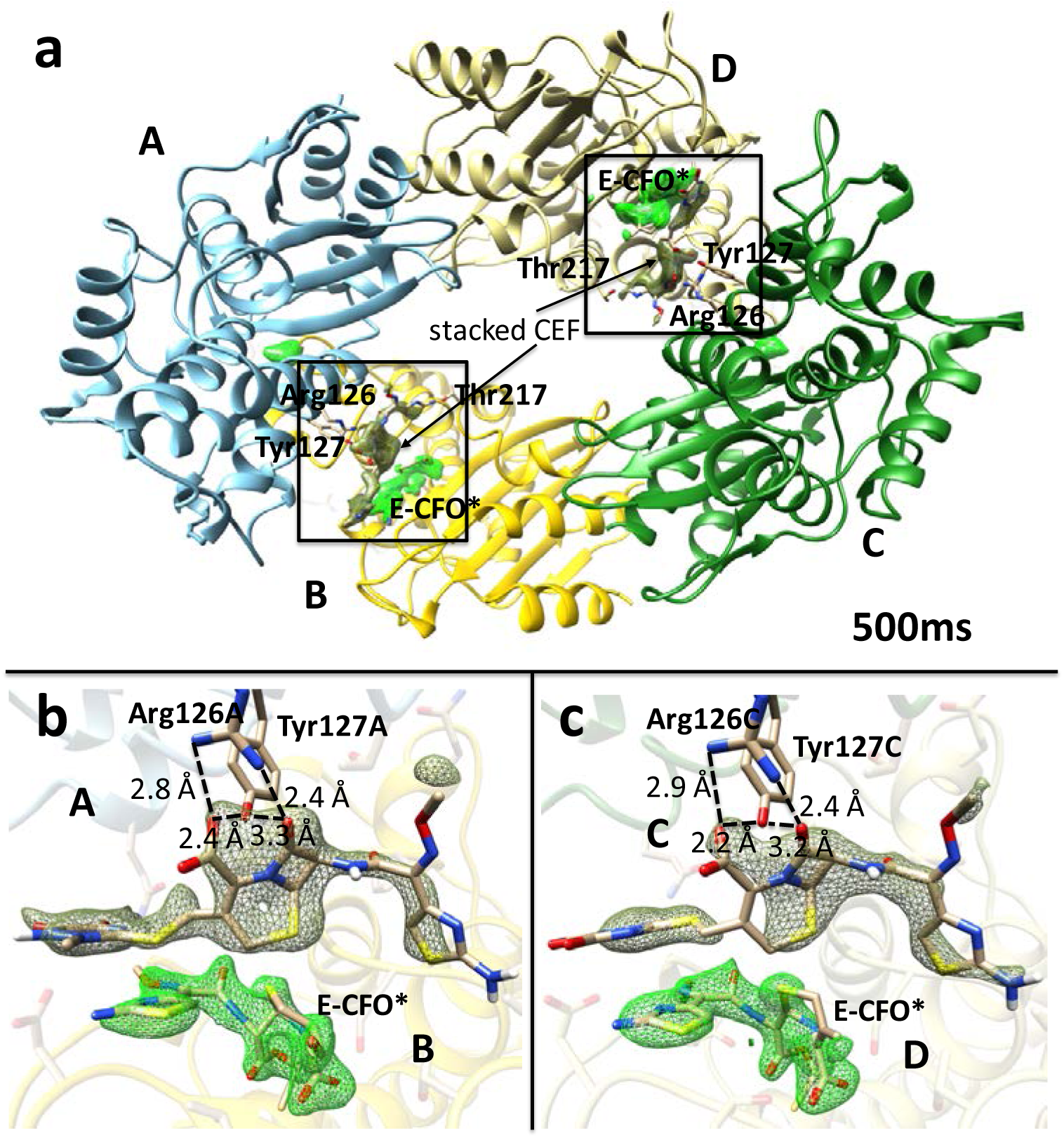
Overview of BlaC as determined in the shard crystal form at 500 ms. The mFo-DFc SA-omit electron density is shown for the covalently bound intermediate E-CFO* in green (contour level: 2.5 σ). Electron density of an additional, stacked ceftriaxone molecule near the active site is shown in dark green (contour level: 2 σ). **(a)** The BlaC subunits A-D displayed in blue, yellow, green and light yellow, respectively. Amino acid residues which interact with the stacked CEF are labeled. Panels **(b)** and **(c)** show enlarged views of the active sites of subunits B and D, respectively. Arg 126 and Tyr 127 with which the respective stacked CEF molecules interact are shown. Some important distances are also displayed (stacked molecules are also observed at the other time delays in the shard crystal form but not in the needles).

**Figure 2.**
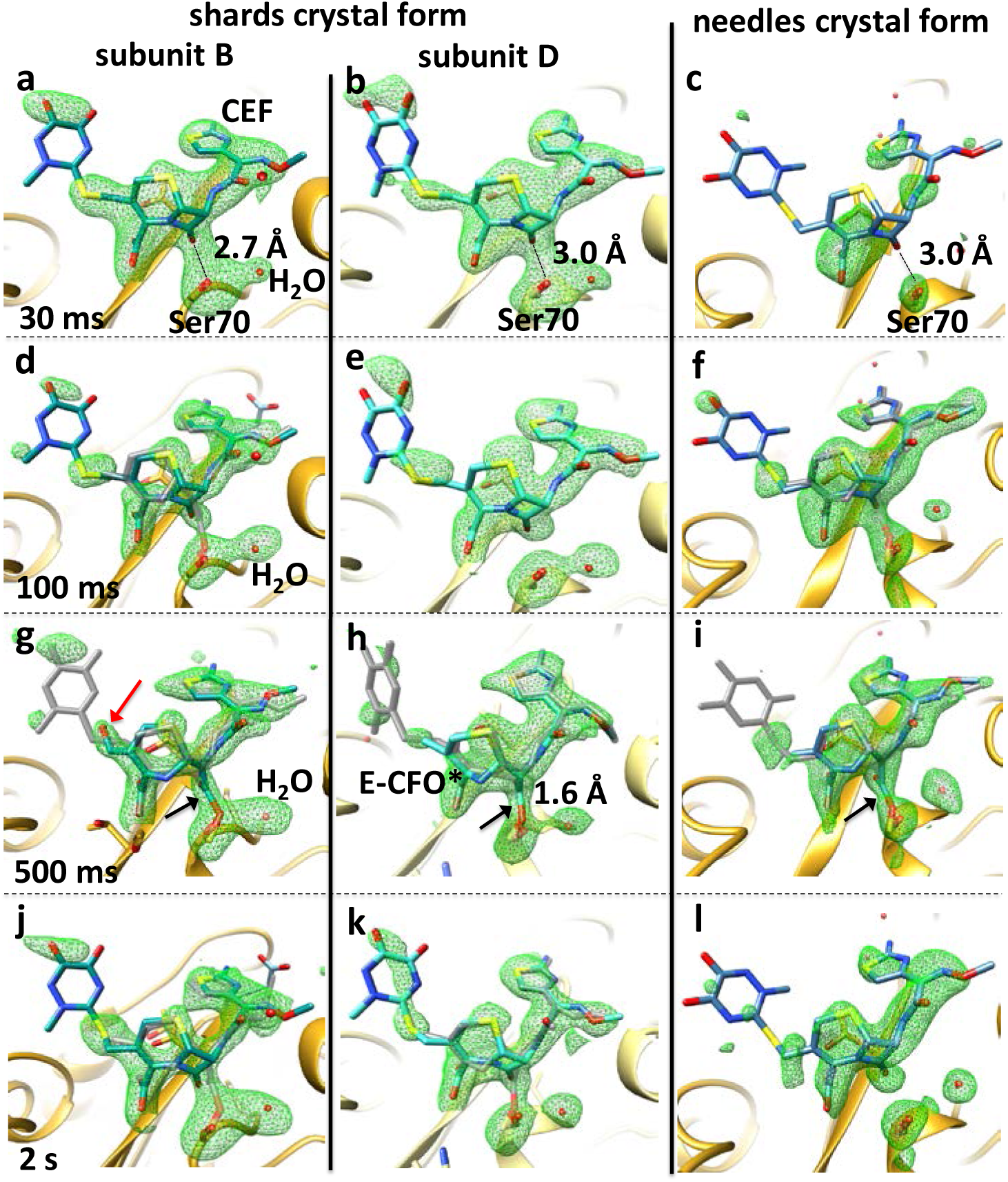
Ceftriaxone electron density in the active site of BlaC in various crystal forms (shards and needles), and at various time delays. The electron density is interpreted by different ceftriaxone species: The main species is displayed in blue, the minor species in gray. For the shard crystal form (panels in the first two columns) mFo-DFc SA-omit density (green) contoured at 2.5 σ is shown. For the needles (panels in the third column), SA omit maps were calculated using extrapolated structure factors (see text) with N=9 for 30ms and 100ms delays, N=6 for the 500ms and N=5 for the 2 s delays. Time-delays are arranged from top (30ms) to bottom (2 s). The first column shows the catalytic cleft in subunit-B of BlaC in the shard crystal form and the second column that of BlaC subunit D. The third column shows the same BlaC region in the monomeric needle crystal from. Black arrows show the electron density of a covalently bound acyl adduct (see Fig. S2 for details). **(a,b,c)** Formation of the ES complex at 30ms. The full length CEF model (blue) is displayed in the active site. The ES complex can be observed at the same time delay regardless whether needles or shards (both subunits) are investigated. **(d,e, f)** Early phases of the formation of a covalently bound CEF adduct observed at 100ms. The full length CEF model (blue) is displayed together with the minor E-CFO* species (gray), where the β-lactam ring is open and attached to Ser-70 in subunit-B (shard crystal form, panel d) and the needle crystal form (panel f). No covalently bound adduct formation is observed in the shard crystal form subunit D (panel e). **(g,h,i)** Fully cleaved and covalently bound adduct (E-CFO* in blue) formation in the active site at 500 ms. A small contamination of full length CEF (gray) is observed. The red arrow points to electron density that may favor the interpretation by an OH group, rather than assuming a double bond. **(j,k,l)** A mixture of the non-covalently bound, full length CEF and covalently bound E-CFO* is observed in the shard crystal form (both subunits) at 2s, while the electron density in the needle crystal form favors only the full length CEF species.

One of the critical questions in MISC concerns whether the enzyme in the crystals is still catalytically active and whether the reaction is limited by constraints of crystal packing or the solvent/precipitant used for crystallization. We have therefore crystallized BlaC in two different crystal forms. With phosphate as precipitant, the BlaC crystallizes in a shard-shaped crystal form with four copies in the asymmetric unit (Fig. 1a) as previously reported (*25*). With PEG 1000 as precipitant, needle shaped crystals are obtained with a monomer in the asymmetric unit (for details see the SM). The packing of BlaC in both crystal forms is shown in Fig. S11. In our MISC experiment, the small microcrystals were rapidly mixed with CEF ‘on the fly’ using highly optimized mixing devices (Fig. S1), and structures of the reaction of BlaC with CEF were determined by scattering from femtosecond X-ray pulses at five time points (unmixed and 30 ms, 100 ms, 500 ms, and 2 s after mixing, respectively) during the reaction in both crystal forms. Results are shown in Figs. 1 and 2. CEF binds to the active site of BlaC as shown in Fig. 1a. In Fig. 2 more details are shown for the substrate binding in the shard and needle crystal forms (see Fig. S2 for details from another viewing direction, and Figs. S3 – S10 for stereo representations of various viewing directions and time points). Strong electron density at 30 ms shows that substrate diffusion into the crystals was successful. At this time delay the formation of the enzyme-substrate complex (ES) is observed (Fig. 2 a,b,c). At 100 ms the ES complex still prevails and is the major component observed. Only on a time scale longer than 100 ms the nucleophilic attack of Ser-70 opens the β-lactam ring. At 500 ms this results in high occupancy of an intermediate which is covalently bound to the enzyme called E-CFO* as shown in Fig. 2 (g,h,i). Since at the same time the leaving group R (scheme 1) is split off, the covalently bound ligand is much shorter than CEF. We see subtle differences between the results from the two crystal forms, and between the subunits in different crystal environments (Fig. 2), confirming previous preliminary observations (*25*). In both crystal forms, at 100 ms a substantial fraction (~70%, see also Tab. S2) of CEF molecules are still intact. A minor fraction (~30%) has an open β-lactam ring (Fig. 2d,e,f). The open, covalently bound species E-CFO* can be identified more clearly at 500 ms, where it dominates the electron density. This confirms, for the first time on a structural basis, previous predictions from biochemical results for other cephalosporin species (*38*). The red arrow in Fig. 2g indicates that the double bond Δ (Scheme 1) may have reacted to an alcohol in subunit B, which does, however, not occur in subunit D, nor in the needle form of the crystals. Additional evidence is presented in the supplemental information, showing the density in the unmixed shard crystal form and a difference map between the 500ms and the 100ms time point showing clear changes in the region of the covalent attachment of the intermediate (Fig. S10c). At 2 s, the structures reveal the steady state of the enzyme, where the binding sites are occupied mainly by the full length CEF with minor contribution from E-CFO* (< 20%) in the shards.

In the multi-copy shard crystal from, subunits A and C do not directly participate in catalysis, at least not in the first 2 s. In the monomeric needle crystal form it appears that the reaction proceeds similarly to that observed in subunit D in the shards. However, substrate occupancy is lower than that in the shards, with substoichiometric occupancy ranging from 20 % - 40 %. The reason for this might be that the enzyme is more tightly packed in the needle crystal form (Fig. S11). To reach full occupancy in the needles, at least 30 mmol/L of CEF (one CEF molecule per asymmetric unit) is initially required, which needs to be delivered by diffusion from the solution to the side of the crystal. While the outside CEF concentration is on the order of 200 mmol/L in both experiments, the ratio of CEF to enzyme varies in the shard and needle crystals. Fig. S11 shows how the solvent volume that contains CEF surrounding the BlaC molecules in the crystals varies. Where it is on the order of 65% for the shards it is substantially lower (38%) in the needles. Fig. S11 also shows that there are substantial differences in the solvent channel sizes in the two crystal forms. Both may significantly impact diffusion of substrate into the crystals. However, compared to other antibiotics such as ampicillin CEF binds relatively slowly to the BlaC (see discussion in the SM). Strong electron density in our maps show that diffusion and formation of the ES complex is near completion on a 30 ms time-scale. Diffusion times and time scales of the ES formation (30 ms) are irrelevant compared the E-CFO* intermediate formation (~500 ms). As a consequence, for our BlaC, the ability to observe the E-CFO* intermediate does not critically depend on CEF diffusion times into the crystals (compare solid and dashed lines in Fig. 3). Accordingly, the reaction dynamics of the catalytic reaction in the needle and the shard crystal forms are similar despite the different crystal morphologies and packing.

**Figure 3.**
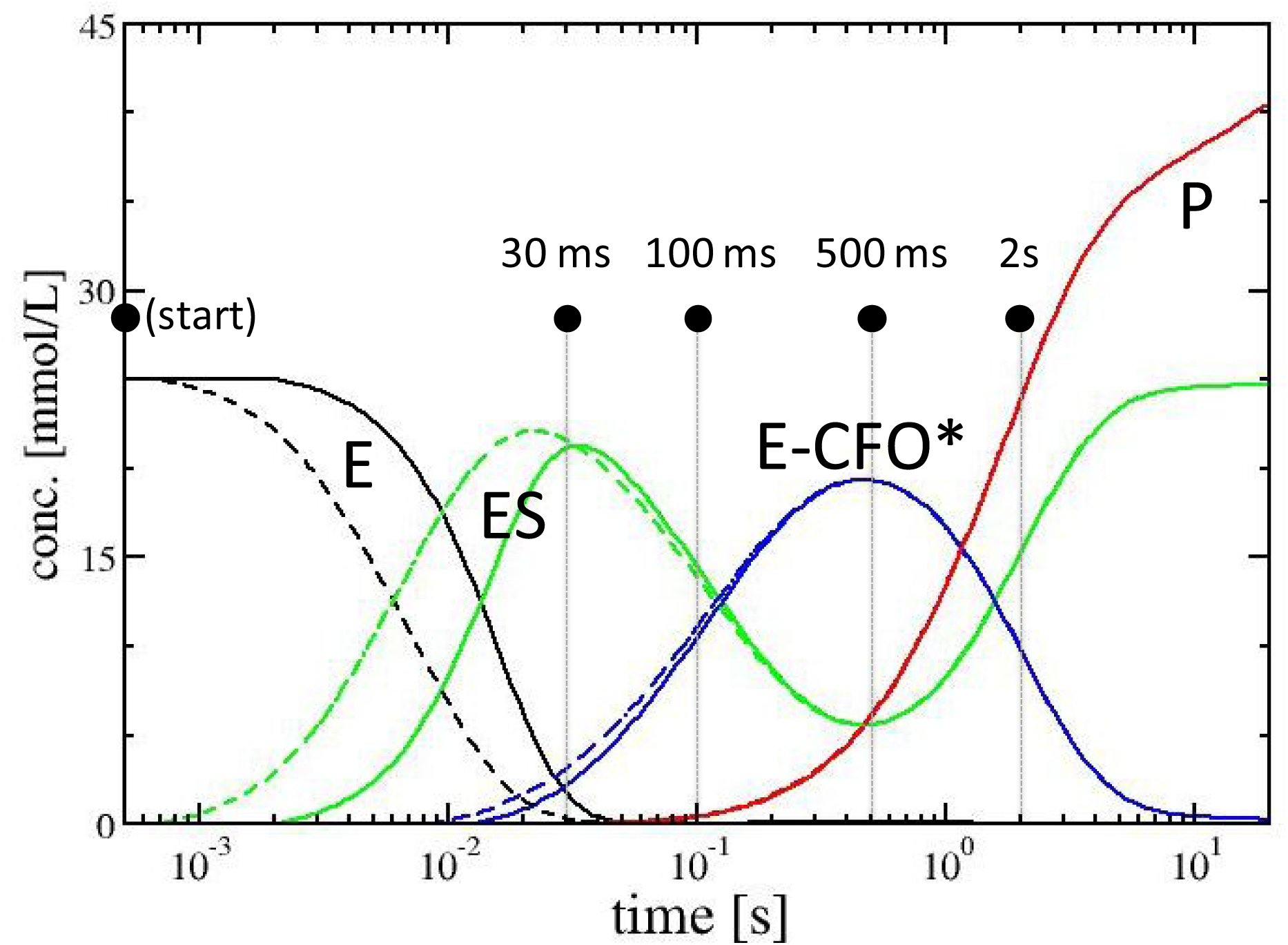
Concentration profile of the catalytic BlaC reaction with CEF as simulated with realistic parameters and a kinetic mechanism as discussed in the SM. The solid lines are calculated with τ_D_ = 15 ms, the dashed lines with τ_D_ = 1.5 ms. Black lines: free enzyme (E). Green lines: enzyme substrate complex (ES). Blue lines: enzyme acyl-intermediate complex (E-CFO*). Red lines: product P (CFO, inactive CEF without leaving group, lactam ring open), released from the enzyme. Black dots: observed time-delays.

An additional CEF molecule (CEF^stack^) can be identified near the catalytic clefts of subunits B and D, each, in the shard crystal form (Fig. 1a,b,c, and Fig. S9). This molecule stacks to the CEF species that occupy the active sites on all time scales. CEF^stack^ is non-covalently interacting with Arg-126 and Tyr-127 of the subunits A or C, which are adjacent to the active catalytic clefts of subunits B or D, respectively (more details are listed in the SM, see also Fig. S9 for a stereo view). This way CEF^stack^ is quite close, pre-oriented, and can rapidly access the active site after the initial CFO has been hydrolyzed and left the enzyme. Since stacking is not observed in the monomeric needle crystal form, it might be argued that it represents a non-physiological, nonspecifically bound substrate that occurs only in dimers in the shard crystals. Previous studies showed that BlaC can crystallize in monomeric form (*37*) as in our needles. Others (*40*) report crystal forms with multiple copies (dimer of dimers) in the asymmetric unit as in our shards. Dynamic light scattering (Fig. S12) and size exclusion chromatography (not shown) performed on the BlaC at pH 5, the same pH as prevalent in our crystallization buffer, show that BlaC can form dimers at this pH, irrespective of the buffer and the concentration. For the BlaC it appears that the oligomerization state in the crystals is determined by the chemical composition of the crystallization buffer. Since, after infection, *M. tuberculosis* resides in the macrophages at pH values near 4.5 (*41*), the BlaC dimer is most likely the physiologically relevant form. The interesting question of the physiological oligomeric state of the BlaC warrants further investigation. The binding of the additional CEF molecule could be an important mechanism to steer the substrate towards, and orient it with respect to, the active site. It appears that at the very high concentrations of CEF applied here, stacking is not required for effective catalysis, as the kinetics in the monomeric needles, where stacking does not occur, is similar to that in the tetrameric shard crystal form. However, when only low CEF concentrations are present, stacking might well be essential to recruit antibiotic substrate molecules to promote effective BlaC function.

One of the major questions addressed here is whether the structural data obtained by MISC can be interpreted in accordance with previous investigations on BlaC catalysis. Ideally, a compatible chemical, kinetic mechanism can developed and expressed in the terminology of enzyme kinetics (*42*, *43*). Accordingly we simulated the catalytic cycle of BlaC using literature values of the Michaelis constant K_m_ (~500 μmol/L) and k_cat_ (0.8 s^−1^) (*36*). We further assumed a sufficiently large number of intermediate states in our mechanism to explain our observations (see text and Tab. S3 in the SM). Our simulations describe the change from the transient state kinetics regime at 30 ms to the steady state regime at 2 s, and covers a large range of substrate and product concentrations. We then compared the simulations to our MISC experiment. Since only 5 time delays are available, the parameters in the mechanism cannot be determined fully quantitatively, but the simulations must reproduce the experimental observations. After initial formation of the enzyme-substrate (ES) complex represented by a non-covalently bound full length CEF, the intermediate E-CFO* has its peak concentration at 500 ms. It has been previously suggested (*37*) that the hydrolytic cleavage of an acyl-adduct from Ser-70 (hydrolysis of species 4 in scheme 1) should be the rate limiting process of BlaC catalysis. Then, the E-CFO* species should be the dominant species in the steady state. However, this is not the case as the ES complex with the non-covalently bound, full length CEF is prevalent (> 70%) in our MISC data at 2 s (Tab. S2). This can be explained by the simulation, if the nucleophilic attack of Ser-70 on species (2) in scheme 1 is inhibited, or slowed down. High product concentrations of > 10 mmol/L are reached already after one catalytic cycle due to the very high enzyme and substrate concentrations (Tab. S3). Unlike in solution, where such product concentrations require minutes to accumulate, high product concentrations are reached after about 1 s. Consequently, product inhibition is plausible. Its structural mechanism, however, remains unknown. In this scenario, after an initial burst over the first second, the nucleophilic attack on the lactam ring by Ser-70 becomes the rate-limiting process (the E-CFO* formation slows down), and the ES complex accumulates in the steady state as observed in our X-ray data. If the product concentration rises even more, even full inhibition of the enzyme is credible. Additional time points would be needed to determine these effects.

Our results show that structural characterization of enzymatically catalyzed reactions on the millisecond time scale is possible. With more conventional X-ray sources, radiation damage prevents the collection of even a single diffraction pattern (*44*) from these small crystals. These difficulties are circumvented by the ultra-short, brilliant hard X-ray pulses available at XFELs. MISC (*25*) can now be employed to investigate a large number of non-cyclic (single pass) reactions in proteins and enzymes, some of which are of immense biological importance, and might be, in addition, important targets for structure based drug design. The reaction kinetics in crystals might well be different in crystal and solution (*45*). However, as long as the crystals are catalytically active, structures of states that are occupied along the catalytic pathway are authentic. With MHz X-ray pulse rates expected at LCLS-II and the European XFEL, multiple, finely spaced time delays may be collected rapidly to allow for a comprehensive description of the biomolecular reaction in terms of structure and kinetics. In the future, with further advances and more XFELs worldwide, MISC will become an important tool to address fundamental questions on enzyme catalysis.

## Acknowledgements

This work was supported the NSF-STC “BioXFEL” through award STC-1231306, and in part by the US Department of Energy, Office of Science, Basic Energy Sciences, under contract DE-SC0002164 (A.O., algorithm design and development) and by the NSF under contract number 1551489 (A.O., underlying analytical models). This material is based upon work supported by the National Science Foundation Graduate Research Fellowship Program to J.L.O. under Grant No. 1450681. The work was also supported by funds from the National Institutes of Health grants R01 GM117342-01 and R01 GM095583, by funds from the Biodesign Center for Applied Structural Discovery at ASU, and the U.S. Department of Energy through Lawrence Livermore National Laboratory under contract DE-AC52-07NA27344. Part of this work was also supported by program-oriented funds of the Helmholtz Association. The structures and diffraction data of the BlaC, unmixed and mixed with Ceftriaxone at 0 s, 30 ms, 100 ms, 500 ms and 2 s for shard and needle crystal forms are deposited to the pdb with the following access codes. Shards: 6B5X (unmixed), 6B5Y (30 ms), 6B68 (100 ms), 6B69 (500 ms), 6B6A (2 s). Needles: 6B6B (unmixed), 6B6C (30 ms), 6B6D (100 ms), 6B6E (500 ms), 6B6F (2s).

## Author Contributions

J.O, J.M.M-G., C.K., D.O., M.D.M., N.N., J.Z., T.N., D.X., E.B., R.F., L.T., G.N.P., Mar.S. prepared and crystallized samples. M.S.H., M. L., J.K., Mat.S. operated the CXI beamline. A.K., G.C., J.K., D.O., M.W., M.H., S.S., S.R-C., J.C., N.N., J.Z., Y.Z., G.N., S.B., U.W., H.N.C., L.P. provided the injector systems, and operated injection at the CXI beamline. S.P., D.O., O.Y., K.P., A.B., T.N., I.P., T.G., V.M., G.S., R.F., P.S., M.F., T.W., N.Z. collected and processed the data. J.O., S.P., J.M.M-G, C.K., D.O., M.H., M.D.M., T.N., I.P., D.X., R.F., P.F., G.N.P., Mar.S. analyzed and interpreted the data. L.T., A.O, Mar.S. conceived the project. P.F., A.O., G.N.P., Mar.S. wrote the manuscript with contributions from all authors.

## Supplemental Material

### Materials and Methods

#### 1. General Overview

Using a continuous-flow mixing apparatus (Fig. S1) we injected active microcrystals of BlaC simultaneously with the β-lactam substrate Ceftriaxone (CEF) into a liquid jet for delivery to the beam as a stream of randomly oriented hydrated nanocrystals *undergoing catalysis*. The catalytic reaction is initiated by solution mixing at the junction of two capillaries(*32*), and the nanocrystals intersected by the X-ray pulse at specific time-points during the reaction. The use of nanocrystals is essential for observation at short times, and for effective and uniform reaction initiation(*23*). The 120 Hz repetition rate of LCLS allowed for the rapid collection of diffraction snapshots at a number of delay times (time-points) after reaction initiation. Accurate reflection intensities were extracted from the snapshots at each time-point by indexing and Monte Carlo type integration (*46*, *47*). By phasing the data using the structural model of the apo-BlaC (PDB-entry 2GDN), we obtained, as a function of time, information on distinct chemical intermediates of β-lactam substrates within the active site of BlaC. The BlaC enzyme requires limited conformational changes to execute catalysis, allowing us to observe the full enzymatic reaction within a crystal.

#### 2. Crystal forms

Cloning, overexpression and purification of *M. tuberculosis* BlaC has been performed as described (*25*). BlaC was crystallized in the shard crystal form as described earlier (*25*). The crystallization conditions were varied slightly to avoid the growth of larger crystals that otherwise need to be crushed to be suitable for ‘mix-and-inject serial crystallography’ (MISC) experiments. An additional crystal form was obtained from a different crystallization condition using the free interface diffusion (FID) method (Kupitz et al, 2014). In a 1.5 mL eppendorf, 250 μL of a precipitant solution (35% PEG 1000, sodium acetate pH 5.0), were slowly added drop wise through 250 μL of a protein solution at 10 mg/mL. Needle-shaped crystals grow at room temperature in about 48 h. The microcrystalline sample was highly monodisperse with crystal sizes between 3 and 5 μm as demonstrated by dynamic light scattering. Microcrystal quality was assessed with SONICC (second order of non-linear imaging of chiral crystals). The suspension showed an intense SONICC signal demonstrating the crystallinity of the sample. X-ray powder diffraction was used as a quality test to verify the existence of diffracting crystals. A very high density pellet of micro-crystals was transferred to a transparent plastic capillary (MITIGEN). A small amount of precipitant solution was kept to prevent crystals from drying out. The capillary was mounted onto a regular goniometer base and data were collected for 3 min on a home source. Intense powder rings were observed up to 7 Å. Weaker rings were also observed to extend up to approximately 4 Å.

#### 3. Injectors

The mixing injectors used in this experiment were based on the design by Calvey et al.(*30*) shown in Fig. S1. In these devices, crystal suspension and a buffer (either 1 mol/L Na-phosphate or Na-acetate, pH 5) containing 200 – 300 mmol/L CEF flow in coaxial capillaries. The flows are combined and forced into a constriction, thinning the crystal flow to a narrow jet and allowing rapid CEF diffusion. By varying the length of the device, the sample and buffer flow rates, or placing an expanded region after the constriction, we were able to probe timescales ranging from 30 ms to 2000 ms. Two HPLC pumps (SHIMADZU LC-20AD) drove the flow. Crystals were held in a custom reservoir built by CXI staff, while buffer was held in a larger reservoir (KNAUER VARILOOP), allowing water flow through the HPLC pump without diluting either sample or buffer. A pressure controller (PROPORTION-AIR GP1) was used to regulate helium pressure in the device. For each condition, the solution is considered mixed when the CEF concentration exceeds 40 mM, sufficiently high to cause rapid binding. The reported mixing times are the time for the concentration around the average crystal to reach this threshold, with upper and lower bounds given for the first and third quartile. In these calculations, the crystals are assumed to be much smaller than the focused jet, and fluctuations in flowrate are neglected. The mixing times for each time point are reported in Tab. S1. The delay time is defined as the time that the reaction is allowed to proceed after mixing. During this time, the crystals traverse the device before being probed by the x-ray beam. Uncertainty in the delay time results from errors in the sample and buffer flow rates (which come from the factory specifications for the Shimadzu LC-20AD HPLC pumps that we used to drive the flows) and from small variations in the diameters and lengths of capillaries used to make the mixing injectors. Mixing injectors were designed so that the delay time slightly exceeded the nominal timepoint to allow for additional time for the ceftriaxone to diffuse into crystals. Tab. S1 lists the delay times and flow parameters for different timepoints.

#### 4. Data collection, data analysis and structure determination

Serial femtosecond crystallography (SFX) experiments were performed at the Coherent X-ray Imaging (CXI) instrument (*48*). Microcrystals were mixed with the antibiotic ceftriaxone (200 – 300 mmol/L) before injection into vacuum using a mixing jet injector described above that allowed millisecond time resolution. Diffraction patterns were recorded on a Cornell-SLAC Pixel Array Detector (CSPAD) operating at 120 Hz to match the X-ray pulse frequency. Data for shards and needles were analyzed in an identical fashion. Cheetah (*49*) was used to filter out diffraction patterns containing Bragg reflections. These patterns were indexed and integrated using the CrystFEL (version 0.6.2) program suite (*47*, *50*). Partial intensities were scaled and merged using linear and Debye Waller factor scale factors. Data statistics is listed in Tab. S2. The BlaC structures were solved for the needles and shards using molecular replacement by PHASER (*51*). For the shards, the tetrameric structure determined by Kupitz et al. (2017) was used as initial model. For the monomeric structure in the needles, subunit D of the tetramer was extracted and used as a search model. Reference structures S_ref,n_ and S_ref,s_ were determined for the <u>n</u>eedles and <u>s</u>hards, using the respective ‘unmixed’ data for both crystal forms. To determined structural changes after mixing, difference maps were determined. For the shards, unit cell changes on the order of 2 Å and larger were observed after mixing. This prevents the calculation of isomorphous difference maps. With the needles however, unit-cell changes were not observed (Tab. S2) and isomorphous difference maps can be calculated. Accordingly, two different strategies were followed to analyze the two types of data.

i. Structures for the shard crystal form: Since isomorphous difference maps could not be calculated, structural interpretation has been based on omit difference maps. The tetrameric reference model was refined using simulated annealing (SA) in ‘PHENIX’ against the observed 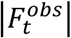. For this refinement water and phosphate molecules residing in the active sites of all subunits were removed. In addition Ser-70 was replaced by a glycine (Gly-70) in subunits B and D. The structure was heated to 5000 K (default) and slowly cooled to 300 K. As a result, a model of the apo-protein without any ligands in the active site was obtained. After the refinement, mFo-DFc omit difference maps 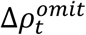 were calculated for each time point t, where the Fo correspond to the 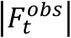 and the Fc are determined from the refined (partial) model, m is the figure of merit and D is a coordinate error dependent weighting term (*52*, *53*). The resulting omit-map is essentially free of phase bias towards the ligand free ‘unmixed’ structure. Strong electron density appeared in subunits B and D that was reminiscent of CEF molecules. In subunits A and C, the electron density of only the phosphate and the water molecules re-appeared, a result that was also previously reported (*25*). Hence, the structures of the catalytic clefts in these subunits A and C were restored back to the reference. The 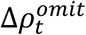 in the catalytic clefts of subunits B and D was exceptionally strong at all time delays (Fig. 2, Fig. S2 – S4, Fig. S6). Appropriate CEF species (Scheme 1) were placed in the positive 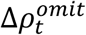 and initially real-space refined in ‘Coot’ using adequate crystallographic information files (CIF), which define the geometry and provide restraints. CIF files were generated for the full length Ceftriaxone (CEF) as well as an open form with the leaving group split off (E-CFO*) as previously described (*25*), compare also Scheme 1. One oxygen of the open lactam carboxyl in E-CFO* was removed and the carboxyl carbon was connected to the Ser70-O_g_ with a weak distance restraint of 1.6 Å. At all time points, either CEF, E-CFO* (bound to Ser-70) or a mixture of both were observed. Their structures were first refined in real space in ‘Coot’. Mixtures of full length, non-covalently bound CEF configurations and Ser-70 bound, open forms (CFO) were refined together in ‘PHENIX’. Note, that E-CFO* was replaced at 500 ms in subunit B with a species displaying an alcohol (Scheme 1d, Fig. 2g, Fig. S6b) instead of the double bond Δ, the structure of which was refined as described. Further refinement including occupancy refinement of the two species was performed with ‘PHENIX’ (*54*) against the 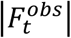. Since a large volume of electron density is shared by CEF and the shorter E-CFO*, occupancy refinement is not reliable. Numbers obtained reflect the fact that the two molecules are present. Essentially complete ligand occupancy is reached at all time delays. Therefore, a potential presence of an unmixed BlaC species was not taken into account during the refinement. The stacked CEF (CEF^stack^) has been added to the last phase of the refinement. The leaving group (the large dioxo-triazin ring) is π–π stacking with the small amino-thiazol ring of the CEF species in the active site resulting in an antiparallel alignment. Distance between the rings are on the order of 3.5 Å. However, as mentioned in the text, the main interactions are with Tyr-127 (between Tyr-127OT_η_ and O_I_ of the CEF^stack^ dihydrothiazine carboxyl) and Arg-126 (between Arg-126N_ε_ and O_I_ of CEF^stack^-O_I_) of the adjacent (non-reactive) dimer subunit (see Fig. 1b,c and Fig. S9). CEF^stack^ is preoriented this way very close to the active site. In order to access the active site, CEF^stack^ only has to flip by 180°, which may be initiated when the CFO species leaves the active site.
ii. Structures for the needle crystal form: Difference structure factor amplitudes 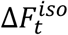 were calculated for each time point t by subtracting the observed reference structure factor amplitudes 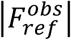 collected with no CEF present from the time-dependent structure factor amplitudes 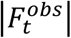. From the 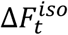 and the phases derived from S_ref,n_, isomorphous difference maps were calculated. In order to model the BlaC structure including (a potentially modified) CEF ligand, conventional electron density maps 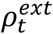 were calculated where the ligand occupancy was extrapolated to 1.0. Extrapolated structure factors 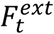 were calculated by adding the 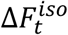 N times (see Tab. S2) to the calculated structure factors derived from S_ref,n_. Extrapolated electron density 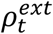 was derived from the 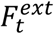. The structures of appropriate CEF derivatives (see above and Scheme 1) were inserted using ‘Coot’ (*55*). At all time points, either CEF, E-CFO* (bound to Ser-70) or a mixture of both were observed (Fig. 2 c,f,i,l). Their structures were first refined in real space against the 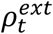 in ‘Coot’. Further occupancy refinement was performed as described above (i).

### Supplementary Text

#### Enzyme Kinetics

The Michealis constant K_m_ is on the order of 500 x 10^−6^ mol/L for BlaC with CEF, and k_cat_ in solution is 0.8 s^−1^ (*36*). The k_off_ rate coefficient of dissociation of substrate from the active site as well as the diffusion coefficient of CEF in the crystals is unknown. Accordingly, we need to assume values that yield plausible results. When the k_off_ rate coefficient is assumed to be equal to the k_cat_ rate coefficient in solution, the k_on_ rate coefficient for the binding of CEF can be estimated from: 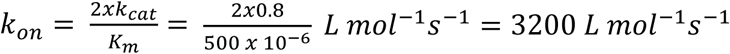. Assuming a diffusion coefficient of 2.3 x 10^−6^ cm^2^/s for CEF in water (*56*) characteristic diffusion times into 2/4/10 x 2/4/10 x 2/4/10 μm^3^ crystals are 0.6ms/2.4 ms/15ms (*23*). The time dependent concentrations of species along the enzymatic pathway were simulated by numerically integrating the coupled differential equations of the simple kinetic mechanism shown in (1) using the above rate coefficients which reproduce the known K_m_. Note, that formation of the covalent E-CFO* complex (acyl intermediate) is irreversible due to the cleavage of the leaving group R from CEF. Tab. S3a lists the parameters that enter the simulation.

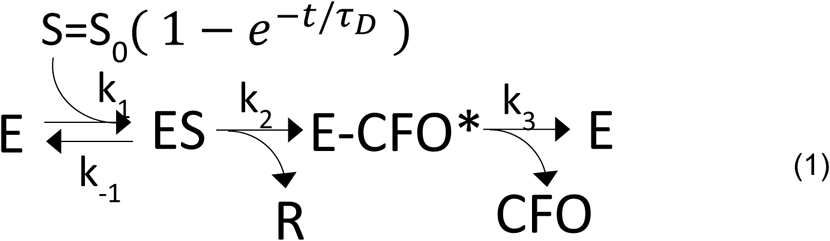

The substrate concentration S in the crystal is determined by the outside substrate concentration S_0_ (50 mmol/L in the simulation) and the characteristic diffusion time τ_D_. S is fed to the free enzyme E and bound to the enzyme with k_on_. The total enzyme concentration was set to 25 mmol/L. Results are shown in Fig. 3 and Tab. 3b for τ_D_ = 15 ms (solid lines) and for τ_D_ = 1.5 ms (dashed lines). Sufficient accumulation (occupancy) of the enzyme substrate complex (ES, green) is achieved after 30 ms even with the longer diffusion time (Fig. 3), which agrees with our observations by MISC (Fig. 2a,b,c). Initially, k_3_ was assumed to account for the rate limiting process and set to k_cat_ = 0.8 s^−1^. As the the low k_3_/K_m_ ratio found in the literature (~ 1.5 x 10^3^ L mol^−1^ s^−1^) suggests, CEF binds slowly to the enzyme. Variations of diffusion times by a factor of 10 (1.5 ms to 15 ms) into the crystals do not play a critical role (Fig. 3, Tab. 3a). The MISC experiment becomes robust against crystal size heterogeneities as well as against mixing time jitter (Tab. S1 and Tab. S3). The ES complex is forming slightly faster with the fast diffusion time, but the kinetics looks essentially the same for both simulated diffusion times. The acyl intermediate (E-CFO^*^, blue) is forming essentially on the same time scale (500 ms) for both crystal forms (Fig. 2 g,h,i). In our MISC X-ray data we do not see clear evidence of a product complex (EP) where the CFO has been hydrolyzed (detached) from Ser-70, and is non-covalently bound to the enzyme. It appears as if this species the enzyme faster than its formation, and therefore does not accumulate sufficiently to be detected. Importantly, the ES complex reappears in our MISC data in the steady state (Fig. 2 j,k,l). That means the E-CFO* cannot be the rate limiting species (with the hydrolysis of the covalent bond the rate limiting process), otherwise it would be E-CFO* which appears in the steady state. However, if ES would initially be the rate limiting species (and the nucleophilic attack of Ser-70 the rate limiting process), E-CFO* would not accumulate sufficiently to be detected so clearly at 500 ms. To solve this dilemma, we assume that rate coefficient k_2_ (the Ser-70 nucleophilic attack) is inhibited by product. Unlike in solution, in the crystal enzyme and substrate concentrations are so high that already after one turnover about 10 mmol/L of substrate is converted to product. Such high free CFO concentrations were never reached in solution studies. Accordingly, on time scales > 1 s, product inhibition of rate coefficient k_2_ was assumed by lowering k_2_ to zero: *k*_2_ = *k*′_2_(1 – *e*^−*P*_*n*_/*I*_*p*_^), with P_n_ the concentration of the released product P divided by an characteristic inhibitory concentration I_p_ in mmol/L. The use of the exponential function is plausible, since the product (Fig. 3, red line) may strongly inhibit catalysis so that the reaction velocity reduces to zero at longer times. This might be the first evidence for product inhibition of the BlaC reaction which awaits further investigations which are outside the scope of this paper. By no means do we suggest that this mechanism is unique. There are only 4 time points (plus the unmixed, free enzyme species). The rate coefficients in the mechanism may vary widely and still reproduce the observations. For instance, the mechanism could be extended by a non-covalently bound CFO (product) which is ready to leave the enzyme. Since we do not see this species, the escape from the protein is most likely much faster than formation of the E-CFO* intermediate resulting in undetectable occupancy. Within a large number of plausible mechanisms, our mechanism is the simplest that explains our experimental observations at limited time points. If a more complex mechanism is to be justified, the collection of time-points is necessary.

**Table S1.** Mixing parameters for each timepoint. The buffer contained 200 – 300 mmol/L CEF in either 1.0 mol/L Na-phosphate (shard crystal form), pH 5, or in 100 mmol/L Na-acetate, pH 5 (needle crystal form).

| Nominal Timepoint (ms) | Mixing Time (ms) | Delay Time (ms) | Sample Flow ($\mu\text{L}/\text{min}$) | Buffer Flow ($\mu\text{L}/\text{min}$) | Constriction Diameter ($\mu\text{m}$) |
| --- | --- | --- | --- | --- | --- |
| 30 | $5^{+6}_{-3}$ | $42 \pm 2$ | $4.0 \pm 0.5$ | $66.0 \pm 0.6$ | $75 \pm 1$ |
| 100 | $10^{+13}_{-8}$ | $114 \pm 4$ | $10.0 \pm 0.5$ | $70.0 \pm 0.7$ | $75 \pm 1$ |
| 500 | $7^{+10}_{-5}$ | $510 \pm 20$ | $8.0 \pm 0.5$ | $32.0 \pm 0.5$ | $50 \pm 1$ |
| 2000 | $15^{+20}_{-11}$ | $2300 \pm 50$ | $10.0 \pm 0.5$ | $45 \pm 0.5$ | $75 \pm 1$ |

**Table S2.**
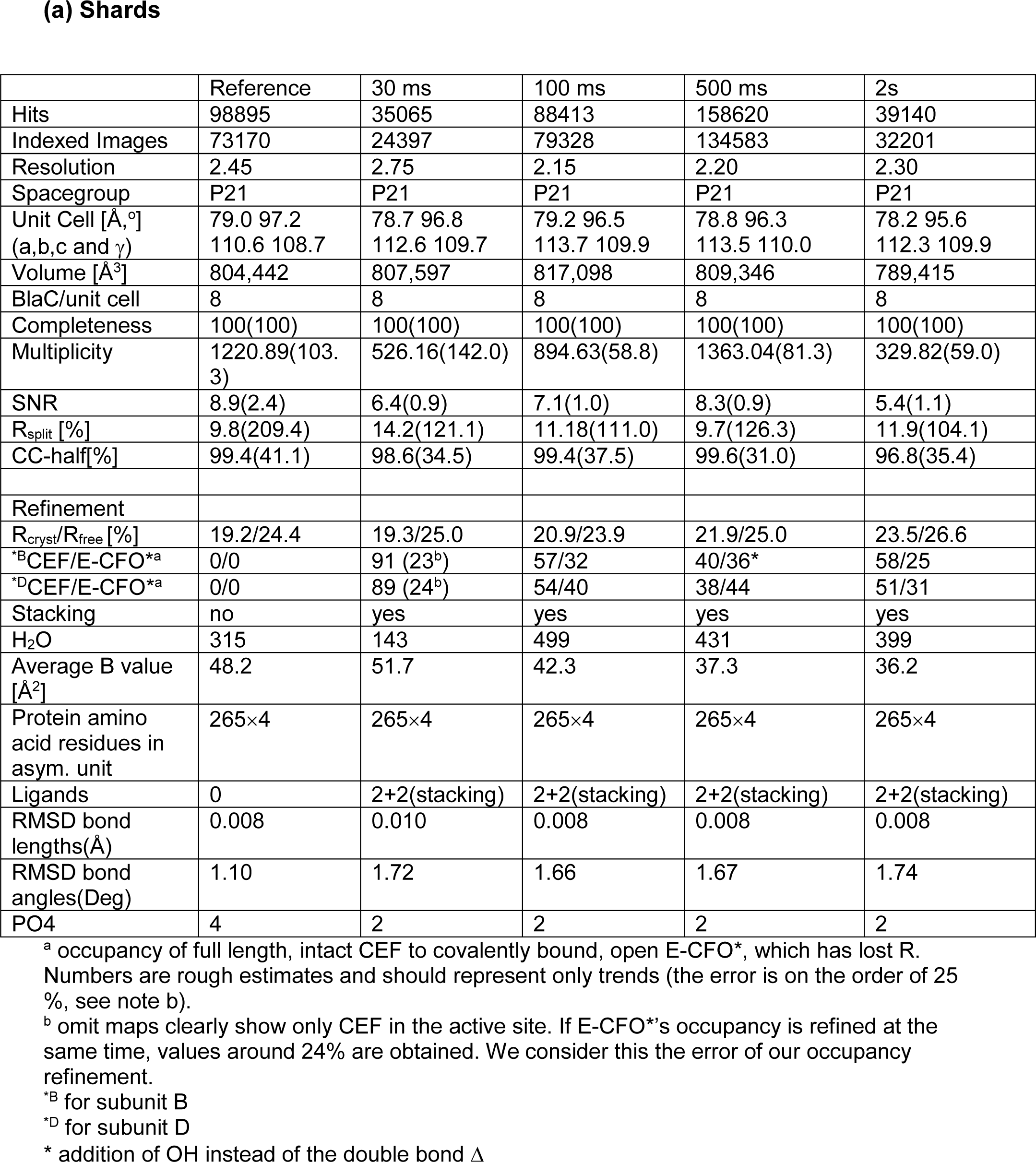
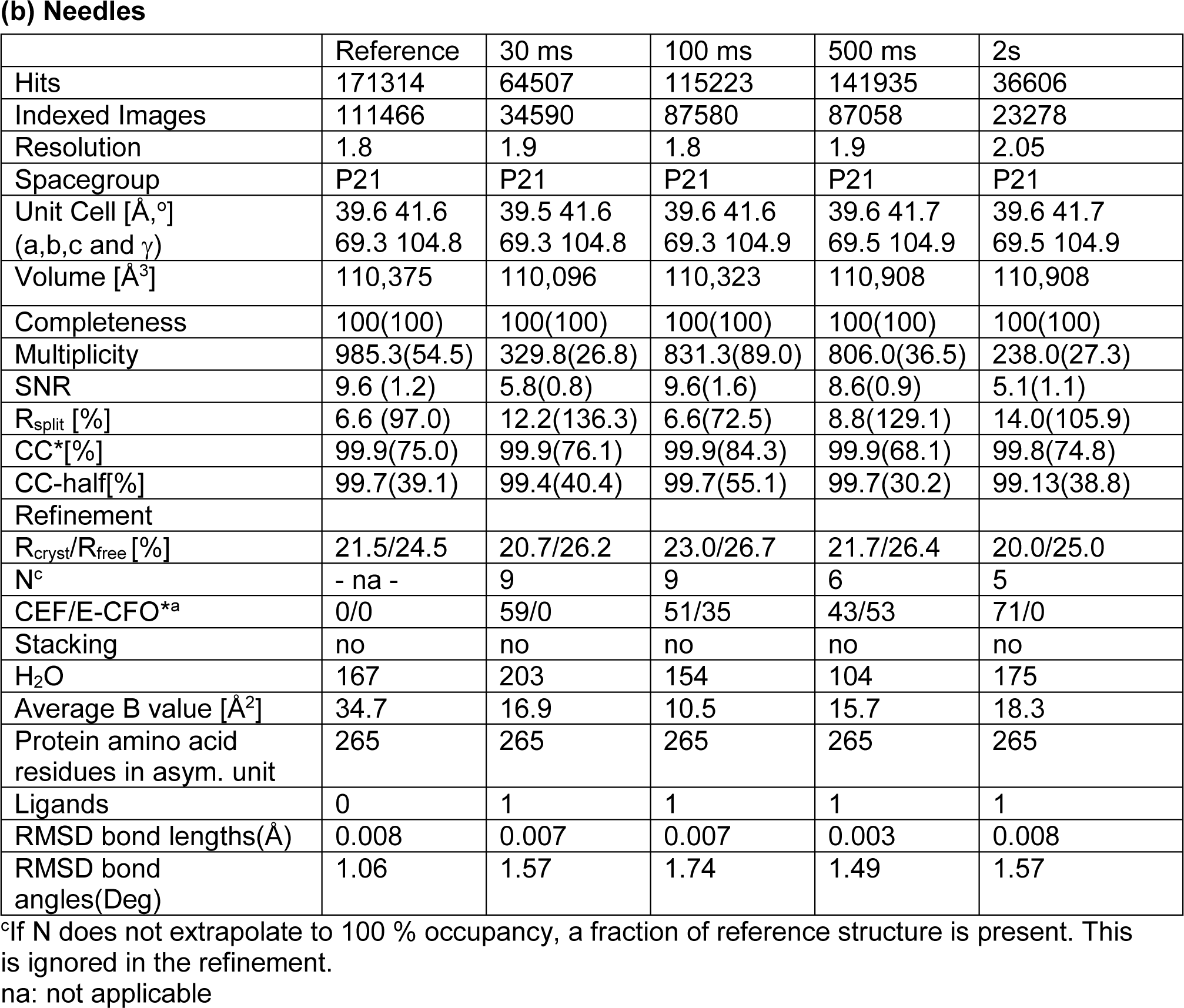
Data collection and refinement statistics.

**Table S3.** Parameter used in the simulation and results. (a) Apparent diffusion time τ_D_ based on crystal size, initial enzyme concentration E_0_, outside substrate concentration S_0_, rate coefficients k_1_ … k_3_, and product inhibition I_p_ (eqn. S1). (b) Occupancy of the various enzyme species as obtained by the simulation. They can be compared to occupancy values listed in Table S2. E: free enzyme, ES: non-covalently bound ceftriaxone in the active site, with leaving group present. E-CFO*: enzyme intermediate with CFO bound covalently, P: free product (CFO).

| $\tau_D$<br>[ms] | $E_0$<br>[mmol/L] | $S_0$<br>[mmol/L] | $k_1$<br>[L mmol<sup>-1</sup> s<sup>-1</sup>] | $k_{-1}$<br>[s<sup>-1</sup>] | $k_2$<br>[s<sup>-1</sup>] | $k_3$<br>[s<sup>-1</sup>] | $I_P$<br>[mmol/L] |
| --- | --- | --- | --- | --- | --- | --- | --- |
| 15/1.5 | 25 | 200 | 3.2 | 0.01 | 7 | 0.8 | 6 |

(b)
|  | 0 ms | 30 ms* | 100 ms | 500 ms | 2s |
| --- | --- | --- | --- | --- | --- |
| E [%] | 100 | 10/2 | 2 | 1 | 1 |
| ES [%] | 0 | 84 | 58 | 19 | 59 |
| E-CFO* [%] | 0 | 6/14 | 40 | 80 | 40 |
| P [mmol/L] | 0 | 0.05 | 0.5 | 5.8 | 23.4 |

\* for 15 ms and 1.5 ms diffusion times, respectively.

**Figure S1.**
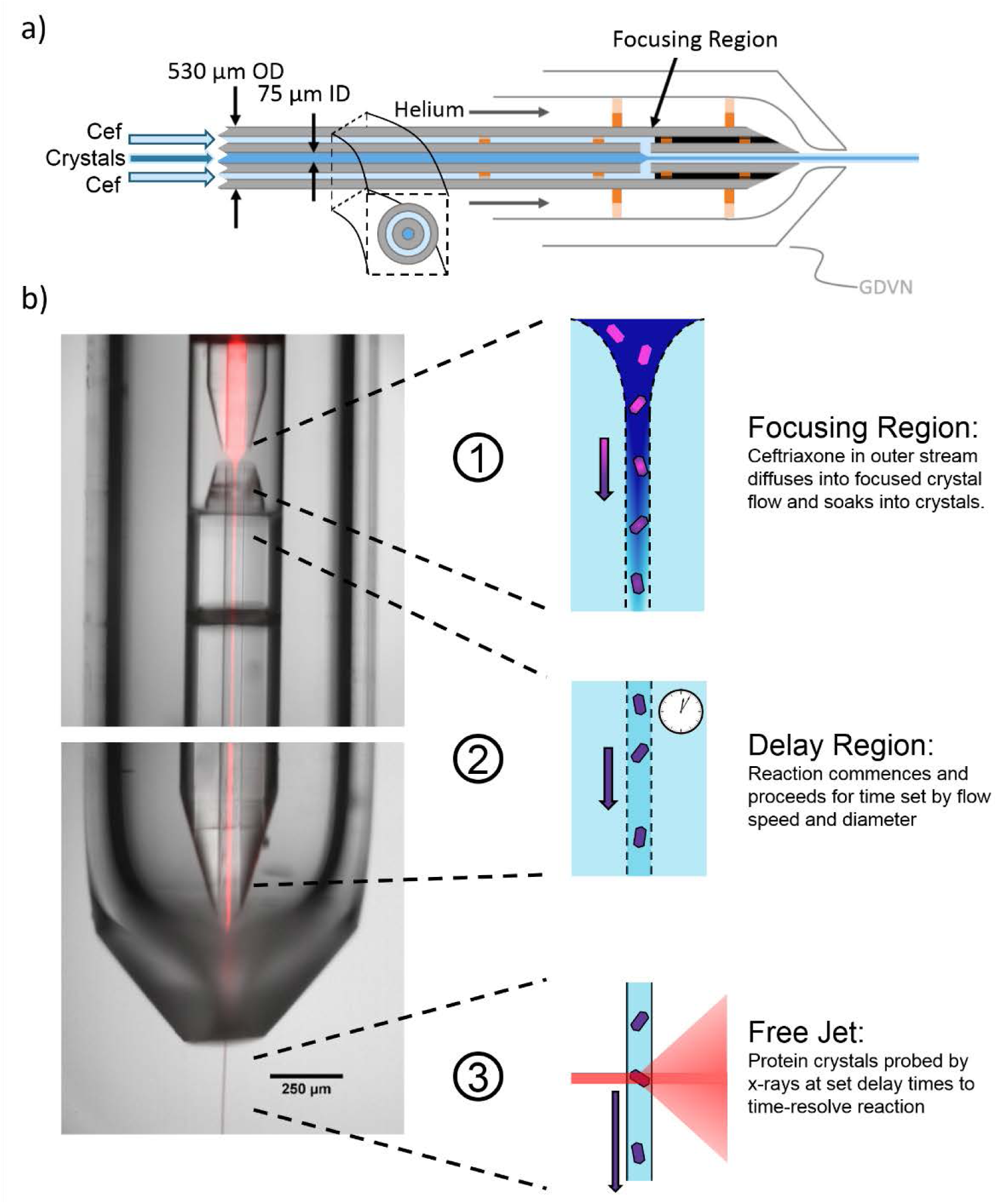
a) Schematic of short timepoint mixing injector. Capillary dimensions vary by timepoint. b) Composite image of fluorescent dye flowing through the sample capillary and water flowing through the buffer capillary. Cartoons illustrate the operating principle of each region of the device.

**Figure S2.**
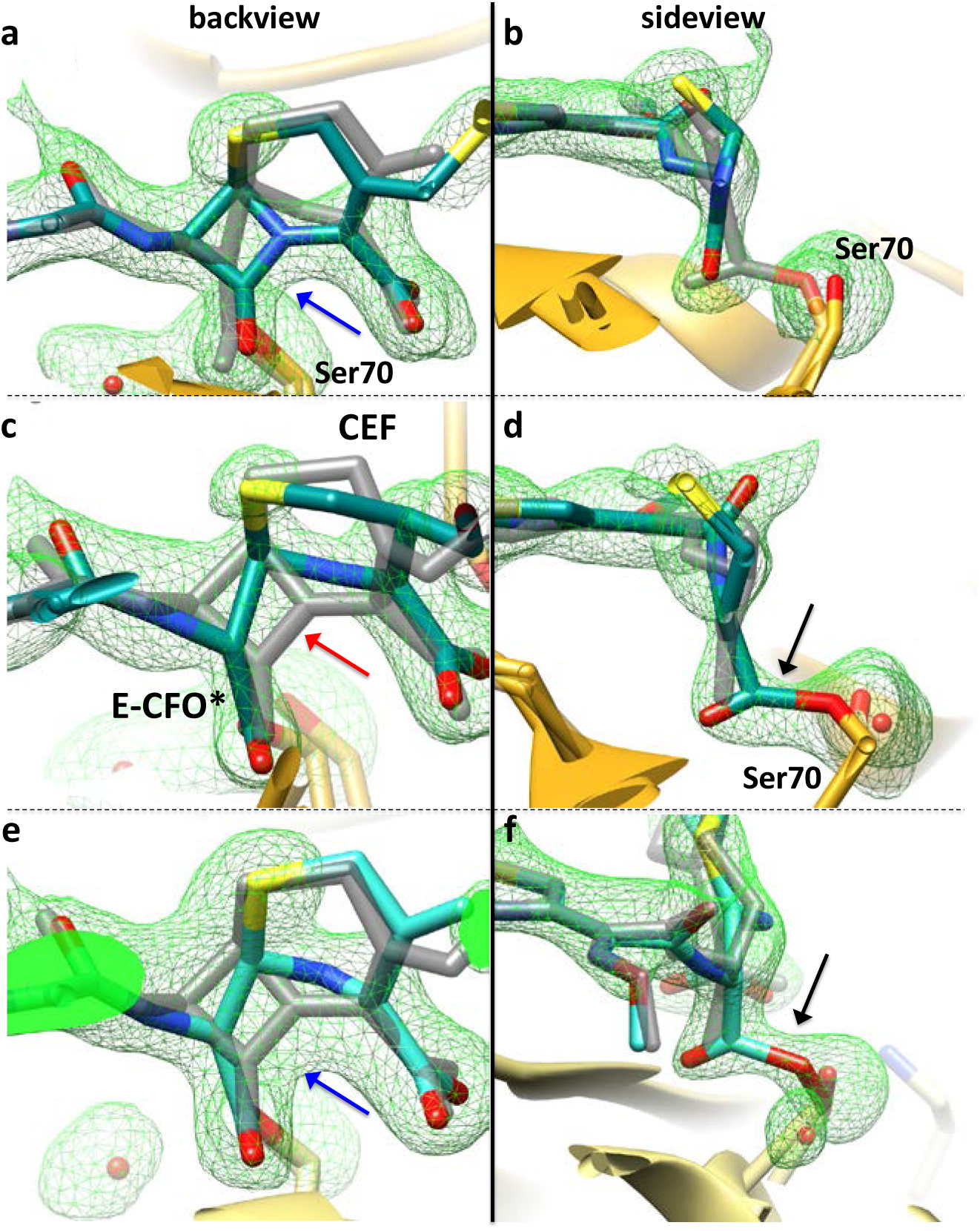
Selected views on the CEF binding site in the BlaC shard crystal form at various time delays. mFo-DFc SA omit electron density (green) contoured at 2.5σ. The first column shows the view on β-lactam ring from the backside in relation to Fig. 1 in the main text. The second column shows the side view to demonstrate cleavage of the lactam ring, and the covalent bond formation to Ser-70. The electron density is interpreted with two species (major species in blue, minor species in gray). **(a, b)** Electron density at 100ms in the BlaC shard crystal form, subunit B. The non-covalently bound, full length CEF is the main species. The closed, uncleaved β-lactam ring nicely fits the electron density (a, blue arrow). The electron density between SER 70 and the open lactam ring is weak (b). The concentration of the covalently bound acyl adduct (E-CFO*) is low. **(c, d)** Electron density at 500ms in the BlaC shard crystal form, subunit B. A covalently bound species (E-CFO*), where the β-lactam ring is opened, and the leaving group is split off, is the main species (blue). **(c)** The closed β-lactam ring poorly fits the electron density (red arrow), and the electron density is interpreted by an open lactam ring (c). Strong electron density between SER 70 and the carboxyl of cleaved lactam ring indicates a covalent bond (d, black arrow). **(e, f)** Electron density at 500ms in the BlaC shard crystal form, subunit D. The full length CEF, and the E-CFO* acyl adduct are present approximately at equal proportions. The β-lactam ring fits nicely in the electron density (blue arrow), which can be interpreted by an uncleaved, full length CEF structure (e). However, in (f), strong electron density between the SER 70 and the cleaved open lactam ring (black arrow) indicates mixing in of a covalently bound E-CFO* species.

**Figure S3.**
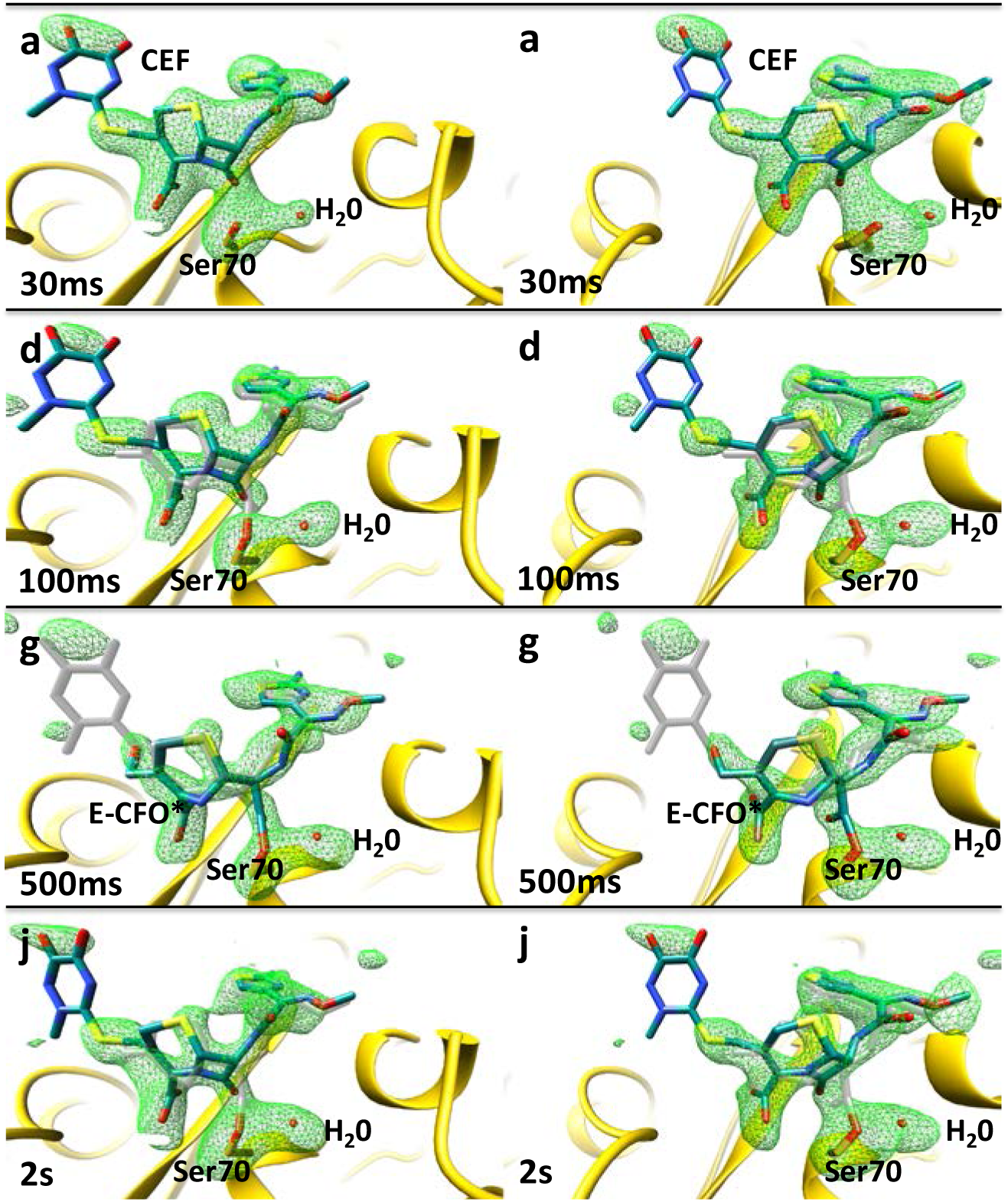
Simulated annealing omit maps, shard crystal form, subunit B (stereo representation of the BlaC reaction with ceftriaxone, from 30 ms to 2 s. Panels are labeled with the same letters as in Fig. 2 (main text). Green: SA-omit difference density (2.5 σ contour). Blue: ligand main structural component, gray: minor structural component.

**Figure S4.**
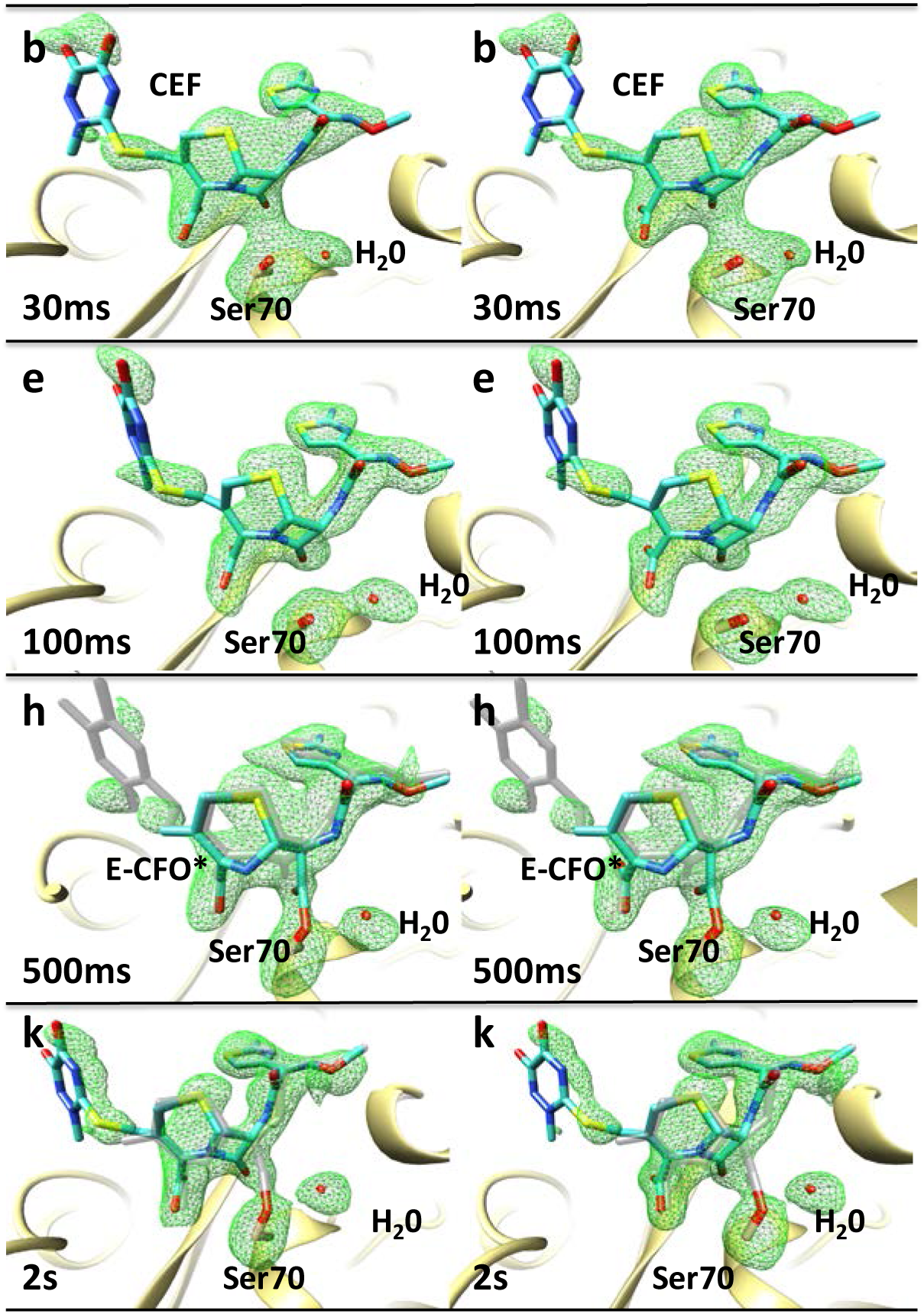
Simulated annealing omit maps, shard crystal form, subunit D. Stereo image of the BlaC reaction with ceftriaxone, from 30 ms to 2 s. Panels are labeled with the same letters as in Fig. 2 (main text). Green: SA-omit difference density on the 2.5 σ contour level. Blue: ligand main structural component, gray: minor structural component.

**Figure S5.**
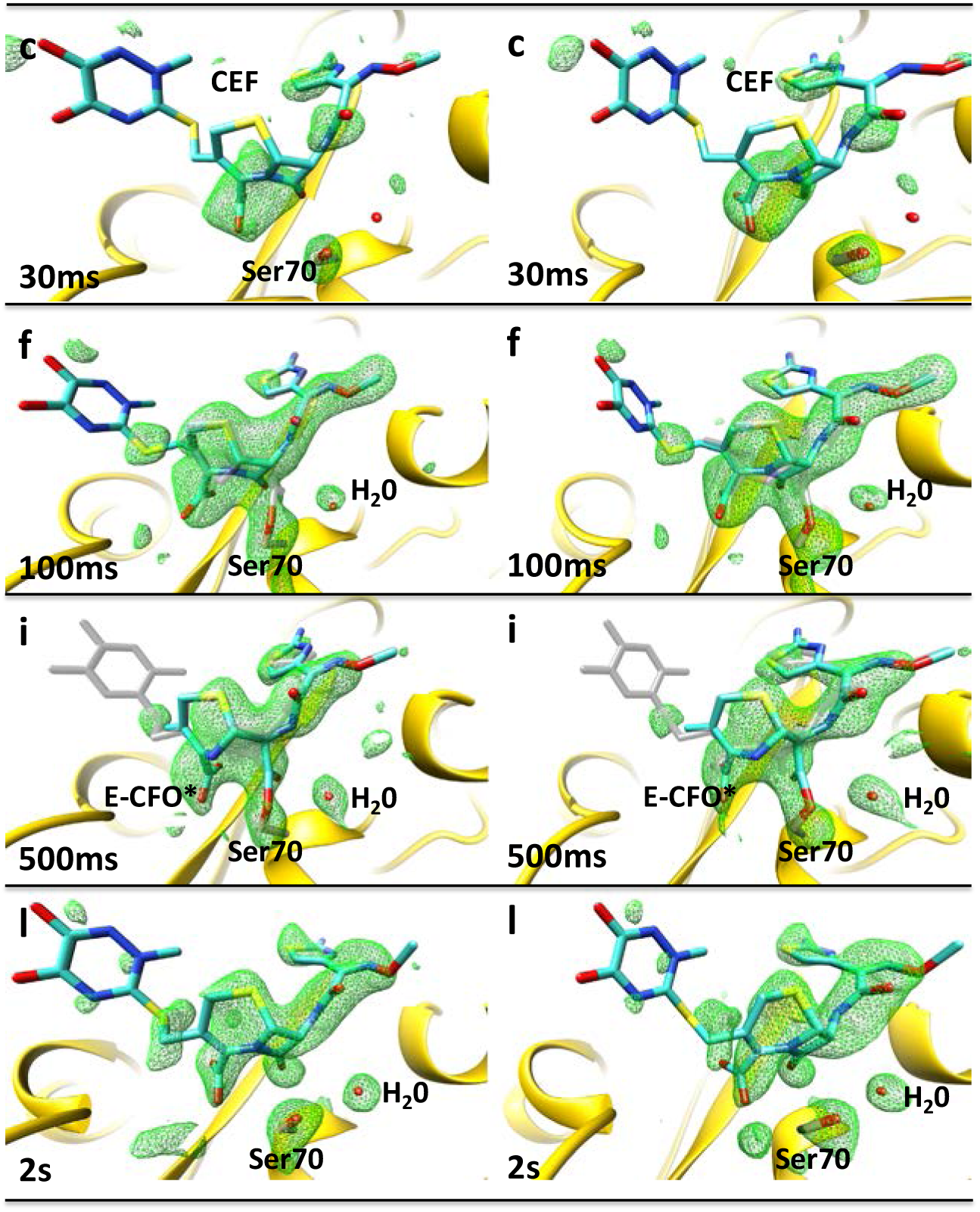
Simulated annealing omit maps, needle crystal form. Stereo images of the BlaC reaction with ceftriaxone, from 30 ms to 2 s. Panels are labeled with the same letters as in Fig. 2 (main text). Green: SA-omit difference density, 2.5 σ contour level. Blue: main ligand component, gray: minor ligand component. The Ser-70 and the nearby water are marked.

**Figure S6.**
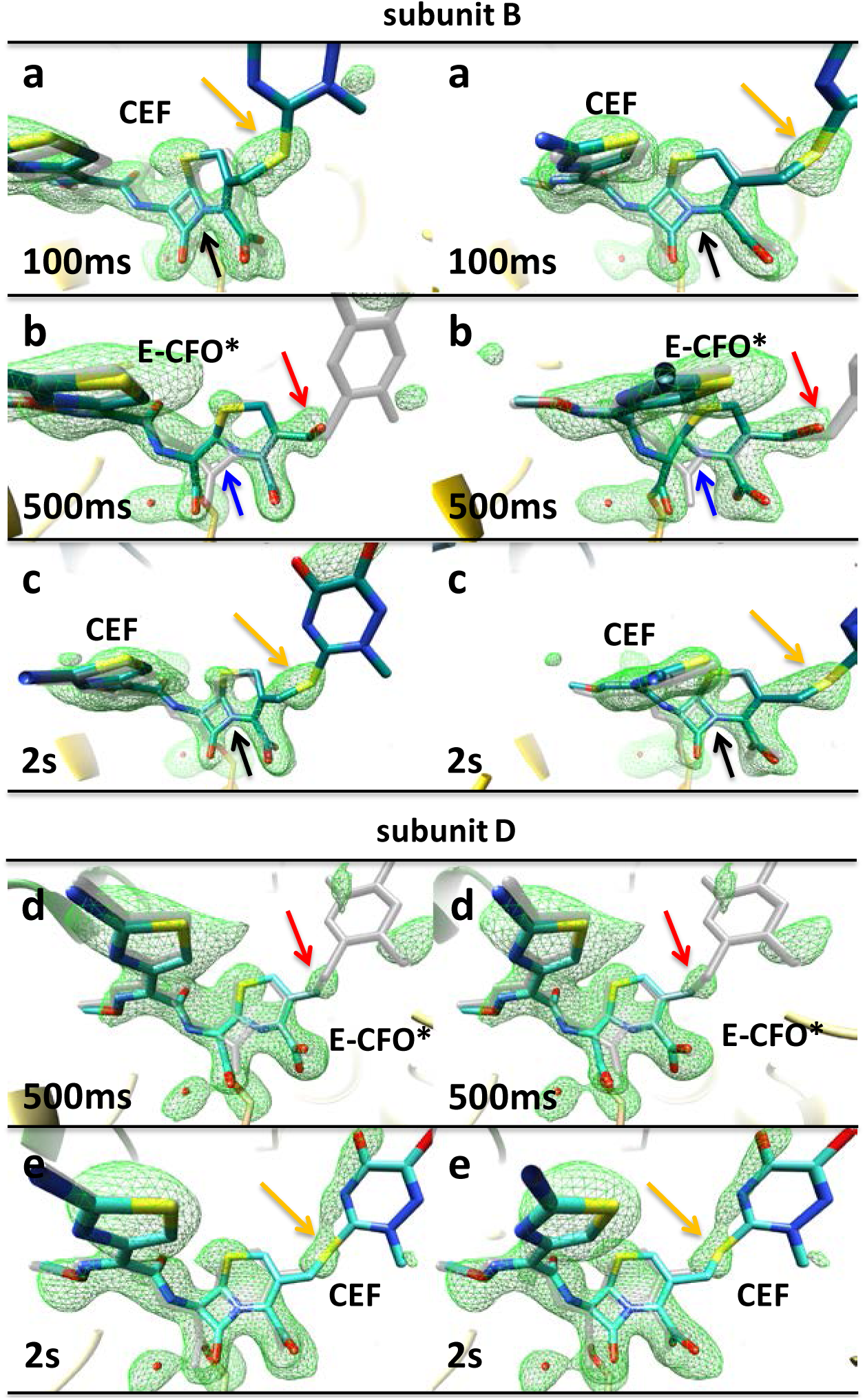
Backside view of ceftriaxone binding to the catalytic cleft of BlaC, subunit B (a,b,c) and D (d,e) of the BlaC shard crystal form at various time delays, mFo-DFc SA omit electron density (green) contoured at 2.5σ. The electron density is interpreted by different ceftriaxone species: main species in blue and minor species in gray. **(a)** Electron density at 100ms (subunit B). Presence of prominent electron density for sulfur (orange arrow) and lactam ring nicely fits the electron density (black arrow) **(b)** Electron density at 500ms (subunit B). Lactam ring is open (blue arrow), the absence of electron density for the sulfur (red arrow) is interpreted as the detachment of R group, followed by the formation of an alcohol. **(c)** Electron density at 2s (subunit B). Presence of prominent electron density for sulfur (orange arrow) and the lactam ring nicely fits the electron density (black arrow). The electron density is interpreted with a full length CEF structure. **(d)** Electron density at 500ms (subunit D). Absence of electron density for the sulfur (red arrow) is interpreted as the detachment of R group. **(e)** Electron density at 2s (subunit D). Reappearance of prominent electron density for the sulfur (orange arrow) and for the dioxo-triaxine ring.

**Figure S7.**
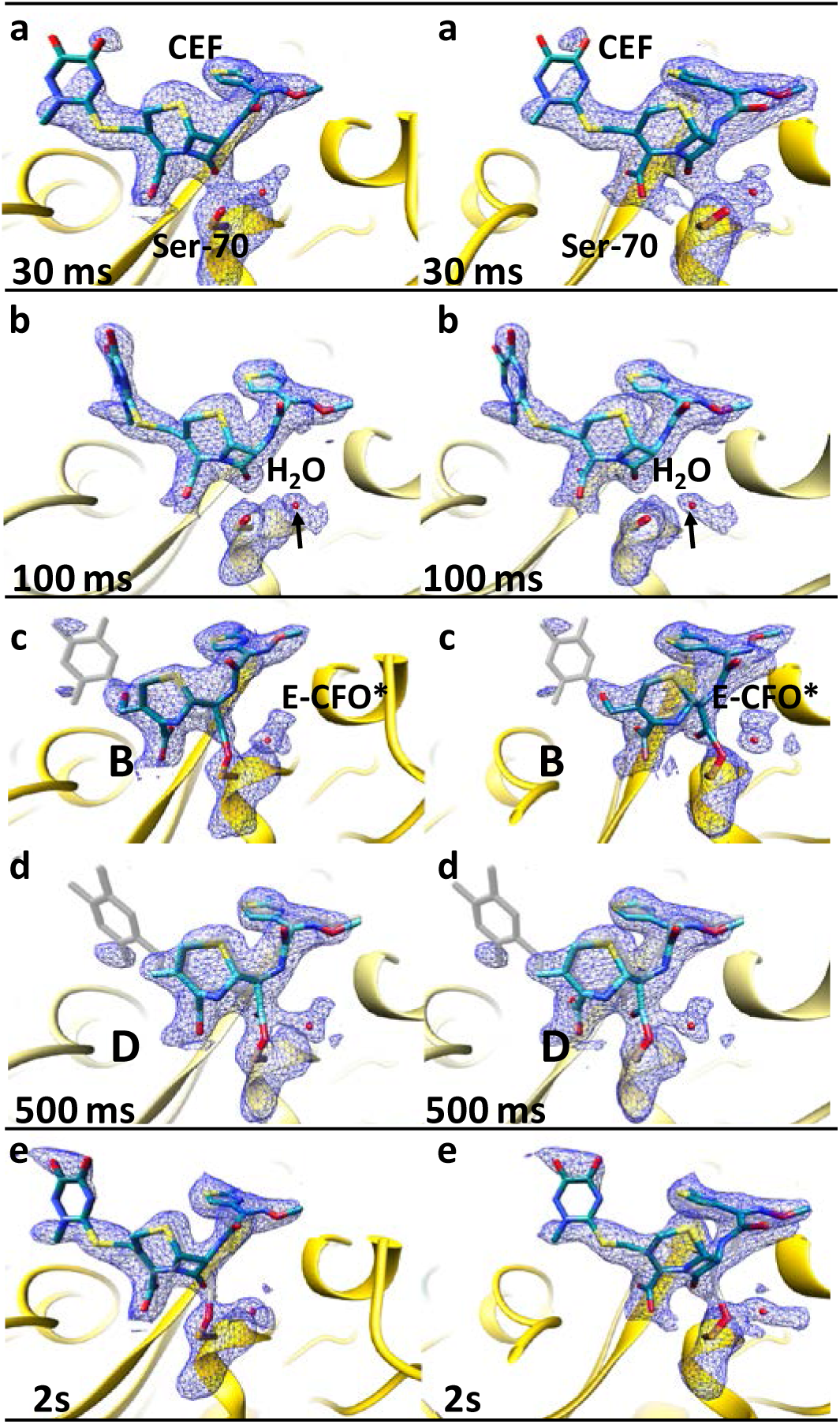
2mFo-DFc electron density (blue, contour 1.1 σ, stereo representation) in the catalytic clefts of subunit B (a,c,e), and subunit D (b,d) of BlaC shard crystal form at different time delay after mixing. The main species is displayed in blue and the minor species in gray. (a) ES complex with the full length CEF non-covalently attached to the active site, (b) mixture of CEF and the covalently bound E-CFO* at 100 ms, (c) and (d) the covalently bound E-CFO* is the main component, (2s) reappearance of the full length CEF with a minor contribution of E-CFO*.

**Figure S8.**
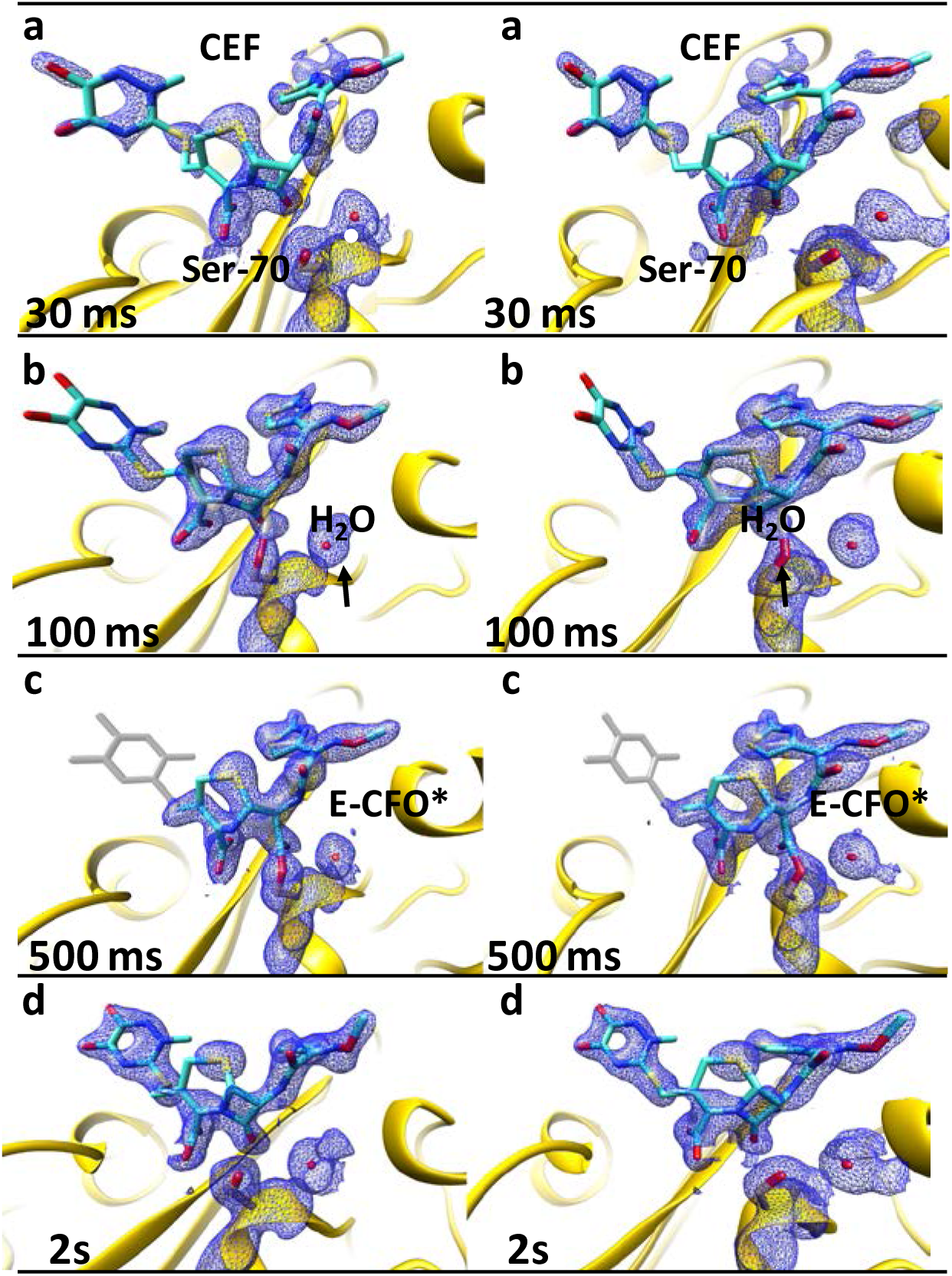
2mFo-DFc electron density (blue, contour 1.1 σ, stereo representation) in the catalytic cleft of the BlaC needle crystal form at various time delays. The maps were calculated using extrapolated structures factors F_o_^ext^ (see text) with N=9 for 30ms and 100ms delays, N=6 for 500ms and N=5 for the 2s delays. The electron density is interpreted by various ceftriaxone species. The main species is displayed in blue and the minor species in gray. **(a)** Formation of ES complex at 30ms. The full length CEF model (blue) is displayed in the active site. **(b)** Early phase of the formation of a covalently bound E-CFO* adduct observed at 100ms. The full length CEF model (blue) is displayed together with the minor E-CFO* species (gray), where the β-lactam ring is open and attached to Ser-70. **(c)** Fully cleaved and covalently bound adduct (E-CFO* in blue) at 500ms. A small admixture of full length CEF (gray) is present. **(d)** 2s, steady state, CEF is dominant species as at 30 ms.

**Figure S9.**
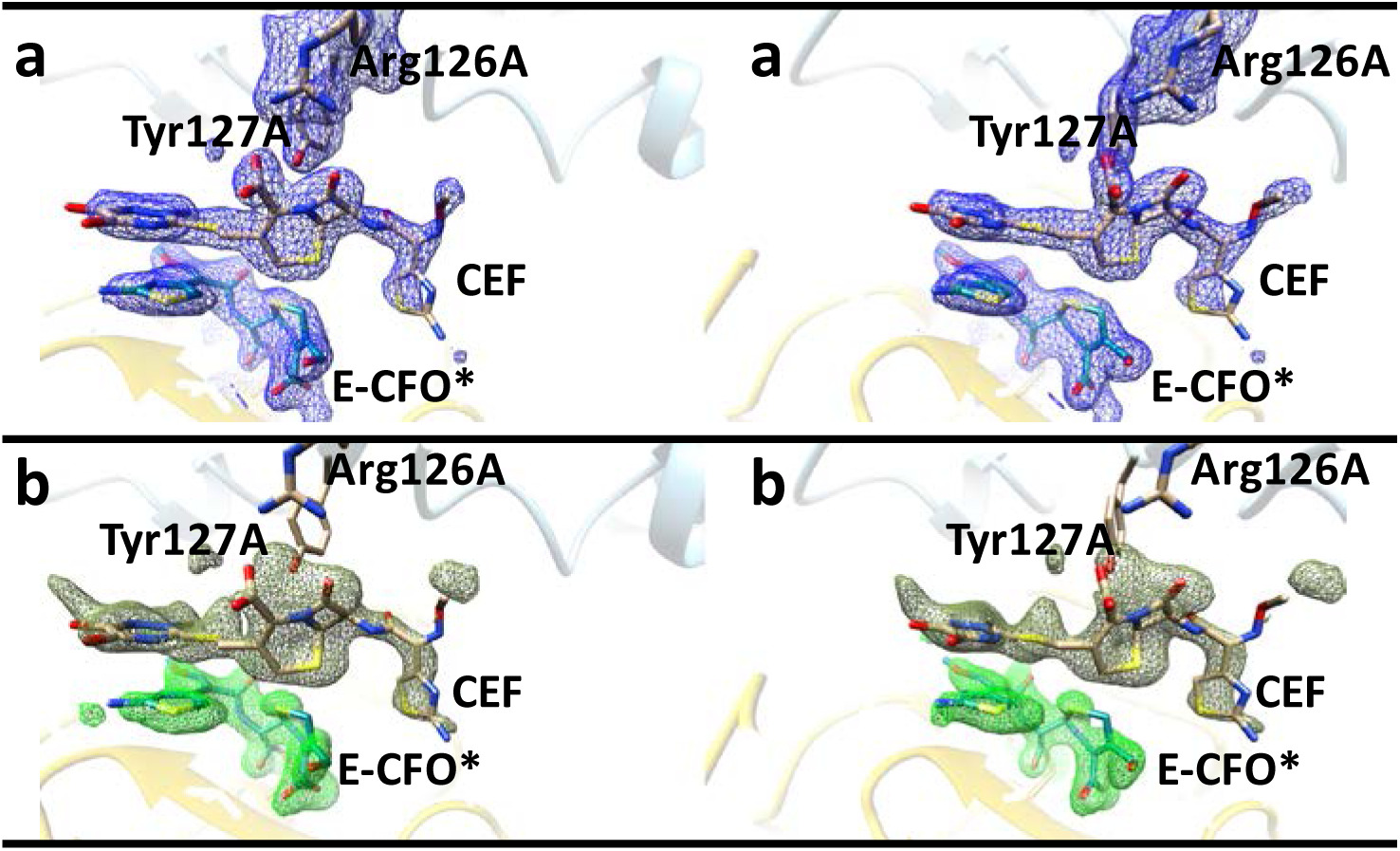
Details in the catalytic cleft of subunit B at 500 ms including the stacked molecule, which interacts with the adjacent subunit A (stereo representation). Details in subunit D with adjacent subunit C are similar. E-CFO*: covalently bound acyl intermediate, CEF: stacked full length ceftriaxone. (a) 2mFo-DFc electron density (1.1 σ contour level). (b) SA-omit maps (2.5 σ contour level for E-CFO*, and 2 σ for the CEF region).

**Figure S10.**
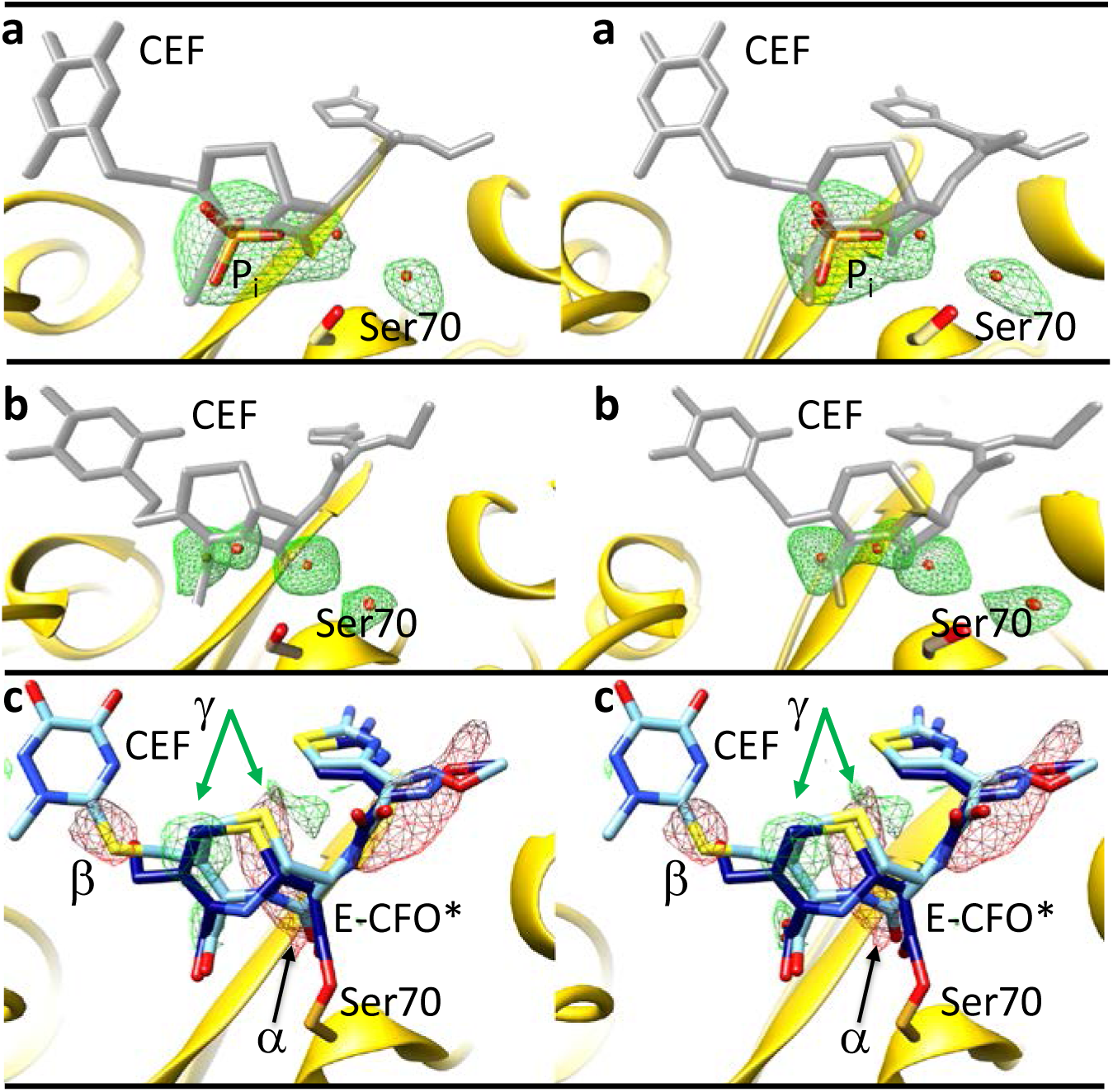
The catalytic cleft of BlaC. **(a)** Unmixed, ligand free structure of the BlaC shard crystals grown in phosphate buffer (stereo representation). The phosphate (P_i_) is marked. The CEF ligand as found at several time delays after mixing is displayed in gray as a guide to the eye. Green electron density: mFo-DFc simulated annealing omit map (2.5 σ contour level) where all small molecules (water and phosphate) in the catalytic cleft were removed. **(b)** Unmixed, ligand free structure of the BlaC needle crystals grown in PEG (stereo representation). The CEF ligand as found at several time delays after mixing is displayed in gray as a reference. Green electron density: mFo-DFc simulated annealing omit map (2.5 σ contour level) where all waters in the catalytic cleft were removed. **(c)** Stereo representation of the F_o_(500ms)-F_o_(100ms) difference electron density map for subunit B of the shard crystals (contour levels: red −2.5 σ, green 2.5 σ). The full length CEF model which is the major species at 100 ms and the covalently bound acyl adduct (E-CFO*) which is the major species at 500 ms are shown in light and dark blue, respectively. The negative feature α (black arrow) is located on the lactam ring carbonyl. This shows that at 500 ms the carbonyl oxygen is displaced, the lactam ring is open, and the covalent adduct has formed. The negative feature β points to a higher sulfur occupancy at 100 ms compared to the 500 ms delay. This is evidence that the leaving group (R) detaches after 100 ms. The positive and negative density pairs on the dihydrothiazine rings (indicated by 2 green arrows) show the shift of the ring positions from back (at 100 ms) to front (at 500 ms) after the lactam ring is opened.

**Figure S11.**
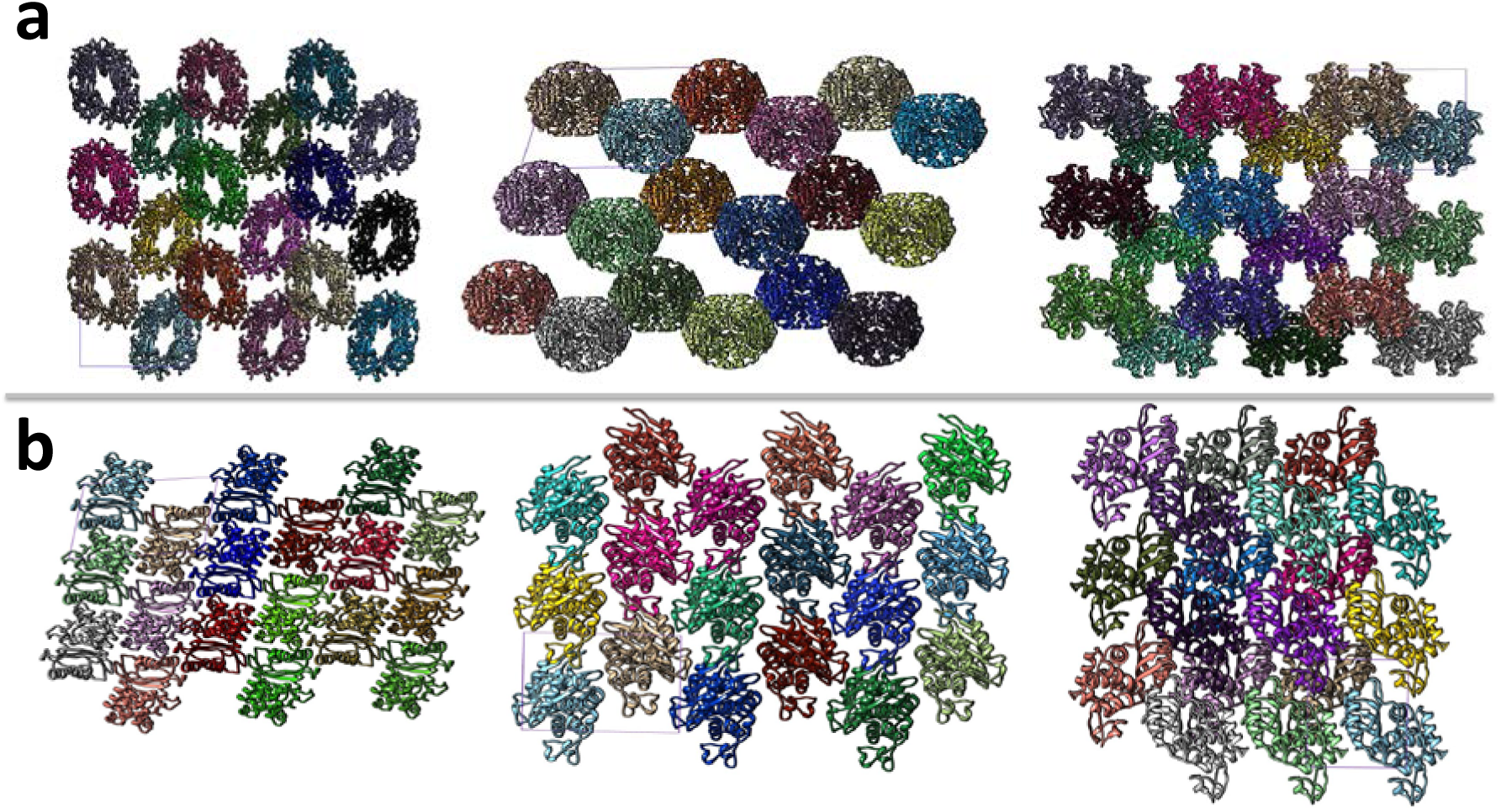
Crystal packing of BlaC in different crystal forms viewed from three different directions normal to the unit cell surfaces. 27 unit cells (three each in the directions along the unit cell axes) are displayed and viewed in orthographic projection. One of the unit cells is outlined for each respective view with faint purple lines. The unit cell volume of the shards is on the order of 805,000 Å^3^ with 8 subunits in the unit cell (one tetramer/asymmetric unit). The concentration of BlaC subunits is 16 mmol/L. The unit cell volume of the needles is about 110,600 Å^3^ with 2 monomers in the unit cell. The concentration of BlaC is 30 mmol/L. (a) Shards, displaying large solvent channels in all 3 directions. (b) Needles, solvent channels are substantially smaller. Note, the display is not to scale. BlaC monomers in (b) appear larger than BlaC subunits in (a).

**Figure S12.**
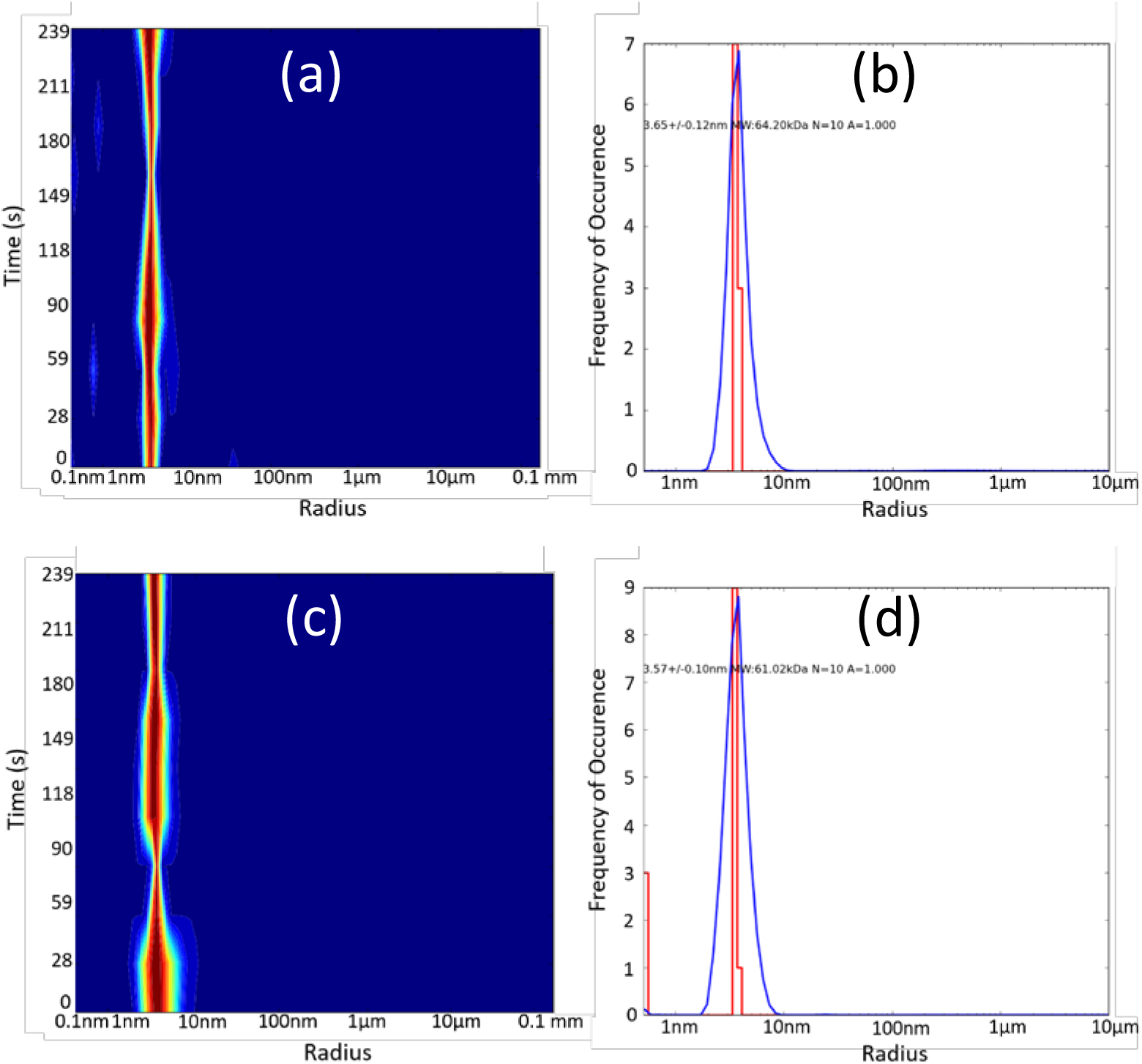
Dynamic Light Scattering on BlaC at 40 mg/mL at pH 5 using a DynaPro NanoStar M3300 (WYATT TECHNOLOGY). A 120 mW laser of 660 nm was used as the light source. For each measurement, the number of acquisitions was 10 and each acquisition time was 20 s. All measurements were carried out at 20 °C. (a) and (b) in 100 mmol/L Na-acetate buffer, (c) and (d) in 100 mmol/L Na-phosphate buffer. (a) and (c) show size distribution over time. (b) and (d) show the radius distribution. A very monodisperse species is present. From (b) and (d) accurate molecular weights can be calculated: (a) 64.2 kDa, (b) 61.0 kDa. The mass of a BlaC momomer is 30.6 kDa (*40*). The BlaC exists as a dimer at this pH in both buffers. Essentially the same result is obtained with 20 mg/mL BlaC.

## Literature

1. C. C. Blake et al., Structure of hen egg-white lysozyme. A three-dimensional Fourier synthesis at 2 Angstrom resolution. Nature 206, 757–761 (1965).

2. L. N. Johnson, D. C. Phillips, Structure of some crystalline lysozyme-inhibitor complexes determined by X-ray analysis at 6 Angstrom resolution. Nature 206, 761–763 (1965).

3. K. Moffat, Time-resolved macromolecular crystallography. Annual review of biophysics and biophysical chemistry 18, 309–332 (1989).

4. M. Schmidt, Structure based enzyme kinetics by time-resolved X-ray crystallography, in: ultrashort laser pulses in medicine and biology. W. Zinth, M. Braun, P. Gilch, Eds., Biological and medical physics, biomedical engineering, ISSN 1618-7210 (Berlin; New York: Springer, c2008, Germany, 2008).

5. M. Schmidt, Time-Resolved Macromolecular Crystallography at Modern X-Ray Sources. Methods Mol Biol 1607, 273–294 (2017).

6. S. Boutet et al., High-resolution protein structure determination by serial femtosecond crystallography. Science 337, 362–364 (2012).

7. G. Chreifi et al., Crystal structure of the pristine peroxidase ferryl center and its relevance to proton-coupled electron transfer. Proceedings of the National Academy of Sciences of the United States of America 113, 1226–1231 (2016).

8. K. Pande et al., Femtosecond Structural Dynamics Drives the Trans/Cis Isomerization in Photoactive Yellow Protein. Science 352, 725–729 (2016).

9. J. Tenboer et al., Time-resolved serial crystallography captures high-resolution intermediates of photoactive yellow protein. Science 346, 1242–1246 (2014).

10. T. R. Barends et al., Direct observation of ultrafast collective motions in CO myoglobin upon ligand dissociation. Science, (2015).

11. C. Kupitz et al., Serial time-resolved crystallography of photosystem II using a femtosecond X-ray laser. Nature 513, 5 (2014).

12. J. Kern et al., Simultaneous femtosecond X-ray spectroscopy and diffraction of photosystem II at room temperature. Science 340, 491–495 (2013).

13. I. D. Young et al., Structure of photosystem II and substrate binding at room temperature. Nature 540, 453–457 (2016).

14. M. Suga et al., Light-induced structural changes and the site of O=O bond formation in PSII caught by XFEL. Nature 543, 131–135 (2017).

15. M. Goelder, R. Givens, Dynamic Studies in Biology: Phototriggers, Photoswitches and Caged Biomolecules. (Wiley VCH, 2005).

16. I. Schlichting et al., Time-Resolved X-Ray Crystallographic Study of the Conformational Change in Ha-Ras P21 Protein on Gtp Hydrolysis. Nature 345, 309–315 (1990).

17. D. Bourgeois, M. Weik, Kinetic protein crystallography: a tool to watch proteins in action. Crystallogr Rev 15, 87–118 (2009).

18. B. L. Stoddard, B. E. Cohen, M. Brubaker, A. D. Mesecar, D. E. Koshland, Jr., Millisecond Laue structures of an enzyme-product complex using photocaged substrate analogs. Nat Struct Biol 5, 891–897 (1998).

19. J. R. Helliwell et al., Time-resolved structures of hydroxymethylbilane synthase (Lys59Gln mutant) as it is loaded with substrate in the crystal determined by Laue diffraction. J Chem Soc Faraday T 94, 2615–2622 (1998).

20. T. H. Kim et al., The role of dimer asymmetry and protomer dynamics in enzyme catalysis. Science 355, (2017).

21. L. A. Sluyterman, M. J. de Graaf, The activity of papain in the crystalline state. Biochimica et biophysica acta 171, 277–287 (1969).

22. S. Geremia, M. Campagnolo, N. Demitri, L. N. Johnson, Simulation of diffusion time of small molecules in protein crystals. Structure 14, 393–400 (2006).

23. M. Schmidt, Mix and Inject, Reaction Initiation by Diffusion for Time-Resolved Macromolecular Crystallography. Advances on Condensed Matter Physics, 1–10 (2013).

24. J. Hajdu et al., Analyzing protein functions in four dimensions. Nat Struct Biol 7, 1006–1012 (2000).

25. C. Kupitz et al., Structural enzymology using X-ray free electron lasers. Struct Dyn 4, 044003 (2017).

26. J. R. Stagno et al., Structures of riboswitch RNA reaction states by mix-and-inject XFEL serial crystallography. Nature 541, 242–246 (2017).

27. H. N. Chapman et al., Femtosecond X-ray protein nanocrystallography. Nature 470, 73–77 (2011).

28. L. Lomb et al., Radiation damage in protein serial femtosecond crystallography using an x-ray free-electron laser. Physical review. B, Condensed matter and materials physics 84, 214111 (2011).

29. H. N. Chapman et al., Femtosecond diffractive imaging with a soft-X-ray free-electron laser. Nat Phys 2, 839–843 (2006).

30. G. D. Calvey, A. M. Katz, C. B. Schaffer, L. Pollack, Mixing injector enables time-resolved crystallography with high hit rate at X-ray free electron lasers. Struct Dyn 3, 054301 (2016).

31. D. Oberthuer et al., Double-flow focused liquid injector for efficient serial femtosecond crystallography. Scientific reports 7, 44628 (2017).

32. D. Wang, U. Weierstall, L. Pollack, J. Spence, Liquid Mixing Jet for XFEL Study of Chemical Kinetics. J. Synchrotron Rad. 21, 1364–1366 (2014).

33. K. F. Kong, L. Schneper, K. Mathee, Beta-lactam antibiotics: from antibiosis to resistance and bacteriology. APMIS 118, 1–36 (2010).

34. M. Bassetti, M. Merelli, C. Temperoni, A. Astilean, New antibiotics for bad bugs: where are we? Ann Clin Microbiol Antimicrob 12, 22 (2013).

35. R. P. Ambler, The structure of beta-lactamases. Philosophical transactions of the Royal Society of London. Series B, Biological sciences 289, 321–331 (1980).

36. J. E. Hugonnet, J. S. Blanchard, Irreversible inhibition of the Mycobacterium tuberculosis beta-lactamase by clavulanate. Biochemistry 46, 11998–12004 (2007).

37. L. W. Tremblay, F. Fan, J. S. Blanchard, Biochemical and structural characterization of Mycobacterium tuberculosis beta-lactamase with the carbapenems ertapenem and doripenem. Biochemistry 49, 3766–3773 (2010).

38. D. B. Boyd, W. H. Lunn, Electronic structures of cephalosporins and penicillins. 9. Departure of a leaving group in cephalosporins. J Med Chem 22, 778–784 (1979).

39. V. Dubee et al., Inactivation of Mycobacterium tuberculosis l, d-transpeptidase LdtMt(1) by carbapenems and cephalosporins. Antimicrobial agents and chemotherapy 56, 4189–4195 (2012).

40. F. Wang, C. Cassidy, J. C. Sacchettini, Crystal structure and activity studies of the Mycobacterium tuberculosis beta-lactamase reveal its critical role in resistance to beta-lactam antibiotics. Antimicrobial agents and chemotherapy 50, 2762–2771 (2006).

41. O. H. Vandal, C. F. Nathan, S. Ehrt, Acid resistance in Mycobacterium tuberculosis. J Bacteriol 191, 4714–4721 (2009).

42. A. Cornish-Bowden, Fundamentals of Enzyme Kinetics. (Wiley-VCH, ed. 4 edition, 2012).

43. J. I. Steinfeld, J. S. Francisco, W. L. Hase, Chemical Kinetics and Dynamics. (Prentience Hall, ed. 2 edition, 1985).

44. J. M. Holton, K. A. Frankel, The minimum crystal size needed for a complete diffraction data set. Acta Crystallogr D 66, 393–408 (2010).

45. S. Yeremenko, I. H. van Stokkum, K. Moffat, K. J. Hellingwerf, Influence of the crystalline state on photoinduced dynamics of photoactive yellow protein studied by ultraviolet-visible transient absorption spectroscopy. Biophysical journal 90, 4224–4235 (2006).

46. R. A. Kirian et al., Structure-factor analysis of femtosecond microdiffraction patterns from protein nanocrystals. Acta crystallographica. Section A, Foundations of crystallography 67, 131–140 (2011).

47. T. A. White et al., CrystFEL: a software suite for snapshot serial crystallography. J Appl Crystallogr 45, 335–341 (2012).

48. S. Boutet, G. J. Williams, The Coherent X-ray Imaging (CXI) instrument at the Linac Coherent Light Source (LCLS). New J Phys 12, (2010).

49. A. Barty et al., Cheetah: software for high-throughput reduction and analysis of serial femtosecond X-ray diffraction data. J Appl Crystallogr 47, 1118–1131 (2014).

50. T. A. White et al., Recent developments in CrystFEL. J Appl Crystallogr 49, 680–689 (2016).

51. R. D. Oeffner, G. Bunkoczi, A. J. McCoy, R. J. Read, Improved estimates of coordinate error for molecular replacement. Acta Crystallogr D 69, 2209–2215 (2013).

52. T. C. Terwilliger et al., Iterative-build OMIT maps: map improvement by iterative model building and refinement without model bias. Acta crystallographica. Section D, Biological crystallography 64, 515–524 (2008).

53. A. Hodel, S. H. Kim, A. T. Brunger, Model Bias in Macromolecular Crystal-Structures. Acta Crystallographica Section A 48, 851–858 (1992).

54. P. D. Adams et al., PHENIX: a comprehensive Python-based system for macromolecular structure solution. Acta crystallographica. Section D, Biological crystallography 66, 213–221 (2010).

55. P. Emsley, B. Lohkamp, W. G. Scott, K. Cowtan, Features and development of Coot. Acta crystallographica. Section D, Biological crystallography 66, 486–501 (2010).

56. M. R. Majidi, K. Asadpour-Zeynali, B. Hafezi, Electrocatalytic oxidation and determination of ceftriaxone sodium antibiotic in pharmaceutical samples on a copper hexacyanoferrate nanostructure. Anal Methods-Uk 3, 646–652 (2011).

